# Bayesian Markov models improve the prediction of binding motifs beyond first order

**DOI:** 10.1101/2020.07.12.197053

**Authors:** Wanwan Ge, Markus Meier, Christian Roth, Johannes Söding

## Abstract

Transcription factors (TFs) regulate gene expression by binding to specific DNA motifs. Accurate models for predicting binding affinities are crucial for quantitatively understanding transcriptional regulation. Motifs are commonly described by position weight matrices, which assume that each position contributes independently to the binding energy. Models that can learn dependencies between positions, for instance, induced by DNA structure preferences, have yielded markedly improved predictions for most TFs on *in vivo* data. However, they are more prone to overfit the data and to learn patterns merely correlated with rather than directly involved in TF binding. We present an improved, faster version of our Bayesian Markov model software, BaMMmotif2. We tested it with state-of-the-art motif discovery tools on a large collection of ChIP-seq and HT-SELEX datasets. BaMMmotif2 models of fifth-order achieved a median false-discovery-rate-averaged recall 13.6% and 12.2% higher than the next best tool on 427 ChIP-seq datasets and 164 HT-SELEX datasets, respectively, while being 8 to 1000 times faster. BaMMmotif2 models showed no signs of overtraining in cross-cell line and cross-platform tests, with similar improvements on the next-best tool. These results demonstrate that dependencies beyond first order clearly improve binding models for most TFs.

## 1 INTRODUCTION

Gene expression is regulated through the binding of transcription factors (TFs) to specific recognition motifs within promoter and enhancer DNA sequences. These binding motifs typically contain 6 to 12 only partially conserved bases [1–3]. Learning quantitative models from experimental data that allow us to accurately predict the binding affinities of TFs to any given sequence is important for quantitatively predicting transcription rates from regulatory sequences.

The task of *de novo* motif discovery is to infer from experimental data a statistical or, equivalently, thermodynamic model that can then predict the binding affinity of a TF of interest for any sequence. Motif models can be inferred from numerous types of experiments [4]. Common *in vivo* techniques are ChIP-seq [5] and bacterial-one-hybrid [6], while most modern *in vitro* approaches are SELEX-based [7–9]. These measurements result in sets of hundreds to millions of bound sequences from which the binding motif model is deduced based on the statistical enrichment of binding sites compared to a background set of unbound sequences or a background model for random sequences.

The dominant model for describing the binding affinity of transcription factors to DNA target sequences has been the position weight matrix (PWM). This model assumes that the binding energy can be decomposed into a sum of contributions from each of the nucleotides in the binding site. By Boltzmann’s law, this is equivalent to assuming statistical independence between nucleotides at different positions of the binding site. The PWM model has been enormously successful, because with its only *3W* parameters for a binding site of *W* nucleotides it achieves high accuracy for predicting the binding affinity of high-affinity binding sites. However, modeling the nucleotide inter-dependency often yields better motif predictions than PWMs [10–12]. One reason is that the stacked, neighboring bases largely determine the physical properties of DNA, such as their equilibrium bending angle, minor groove width, propeller twist, or helical twist. The information on the geometric orientation of the bases propagates within the DNA for several positions before fading out, creating a dependence of the DNA physical properties on nucleotide pairs, triplets and longer *k*-mers. Since TFs recognize their target sites not only using hydrogen bonds but also using their structural fit, TF binding motifs show preferences depending on *k*-mer words [13], particularly in the flanking regions outside the hydrogen bonding core region [14]. Furthermore, alternative binding modes of TFs [15, 16] can lead to poor performance of PWMs.

During the past decade, it has become increasingly evident that weak binding sites in enhancers and promoters play an important role in determining transcriptional activity [17–21], and PWMs have limitations to describe the affinities for weak binding sites accurately. Therefore, various more refined models have been developed that depart from the simplifying assumption of independence of motif positions [22–24]. Prime among them are inhomogeneous Markov models of order *k*, in which the probability to observe a certain nucleotide at position *i* depends on the previous *k* nucleotides at *i − k* to *i* − 1. A zeroth-order Markov model is therefore equivalent to a PWM. Dinucleotide weight matrices (DWMs) are equivalent to first-order models, in which the probability of a nucleotide depends on its direct predecessor, and they have shown improved accuracy over PWMs [25–27].

For Markov models of higher order *k*, the large number of *W* × (4^*k*+1^ − 1) parameters can lead to overfitting on the training data and hence bad predictive performance. To address this limitation, our group had proposed a special type of Markov model, the Bayesian Markov model (BaMM) [28], in which the probability for a nucleotide at position *i* of the motif, for example the last nucleotide in ACTCG, is estimated by adding to the actual counts of ACTCG pseudo counts based on how often the shorter (*k* − 1)-mer CTCG has been observed in the binding sites. The probability for CTCG in turn is estimated by adding its counts to pseudo counts based on how often the word of length *k* − 1, TCG, has been observed, and so forth. This procedure can be derived formally in a Bayesian framework with Dirichlet priors. Our software BaMMmotif indeed improved on previous PWM-based methods for *de novo* motif discovery and binding site prediction on *in vivo* data [28].

Here we present BaMMmotif2, an open-source software written entirely from scratch in C++. It contains a novel algorithm for its seed finding stage, which gives it greatly improved speed and slightly improved sensitivity in comparison to BaMMmotif. We improved the robustness of the BaMM-based motif refinement stage using sequence masking. BaMMmotif2 can also learn positional preference profiles for binding site locations from the training data.

Higher-order models have the ability to learn several low-order motifs overlaid on top of each other [29]. It was therefore surmised that at least a part of the improvements of higher-order models on cross-validation benchmarks using ChIP-seq sequences could stem from learning not only the main binding motif of the ChIPped factor but also, overlaid, the binding motifs of cooperating factors whose binding sites tended to co-occur with it [18]. This would of course defeat the purpose of learning the binding affinity of the ChIPped factor. In a different cell type, for instance, in which different co-binding factors are expressed, such a mixed motif might perform badly. It has also been suggested that more complex models could learn complex, nonspecific sequence biases characteristic of the measurement technique, which would allow them to be distinguished from the background sequences. These platform-dependent biases could result from the library preparation, amplification, and ligation biases [30].

We therefore designed a set of benchmark experiments with a focus on detecting such overfitting (Fig. 1): (I) 5-fold cross-validation on ChlP-seq and HT-SELEX data; (II) crosscell-line validation on ChIP-seq data for the same TFs; (III) model training on ChIP-seq data and testing on HT-SELEX data for the same TFs and (IV) vice versa. Scheme (I) examines how the models generalize to unseen data, especially when data is limited.

**Figure 1.**
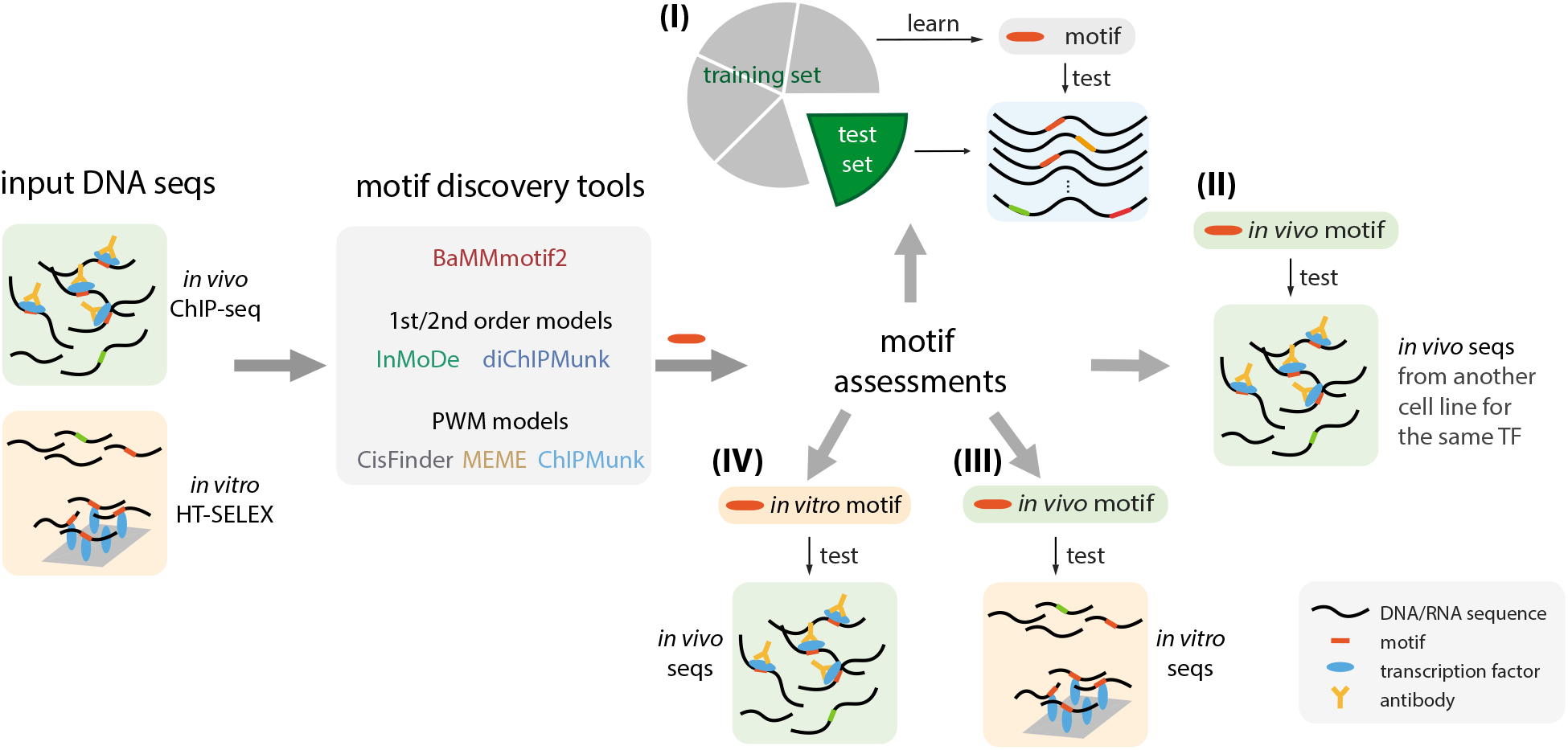
Benchmark pipeline for *de novo* motif discovery. Five state-of-the-art motif discovery tools and BaMMmotif2 learned motif models on *in vivo* and *in vitro* transcription factor binding datasets. The learned models were then assessed (I) by 5-fold cross-validation on the same type of data, (II) by cross-cell-line validation, and (III, IV) by cross-platform validations.

Our results demonstrate that BaMMmotif2 does not show signs of overfitting but rather learns the binding affinity of only the factor of interest, and that BaMMmotif2 is the most sensitive and fastest tool among the ones tested here. Furthermore, BaMMmotif2 keeps improving the performance with increasing model orders and scales better with larger datasets. Finally, we demonstrate the power of BaMMmotif2 models by predicting 1.8 million human CTCF binding sites genome-wide with a false discovery rate below 0.5%.

## 2 MATERIALS AND METHODS

### The BaMMmotif2 algorithm

BaMMmotif2 consists of a seeding stage and a motif refinement stage. The purpose of the seeding stage is to exhaustively identify motifs enriched in the input sequences in comparison to a second-order Markov background model trained also on the input sequences. Each of the motifs below a P-value cut-off is refined by the BaMM-based refinement stage.

#### The fast seeding stage

This method is described in detail in the Supplementary Section 1.1. Briefly, we first count the number of occurrences of each non-degenerate *W*-mer word in {A, C, G, T}^*W*^ (*W* = 8 in this study) in the input sequences. From here on, we only inspect the count array and not the sequences anymore, making the runtime of the seeding stage almost independent of the input set size. By default, reverse complements are mapped to the alphabetically lower of the two *W*-mers.

For each *W*-mer, an enrichment *z*-value is calculated, which is the number of standard deviations with which the observed *W*-mer count surpasses its expected count. The expected count is calculated using a second-order homogeneous Markov model as a background sequence

Here, the hyper-parameter *α_k_* determines how much weight to give to the lower-order. The probabilities of order *k* − 1 are again obtained by adding to the *k*-mer count the pseudo counts from order *k* − 2, and so on down to order 0. In this way, when the number of occurrences observed for (*k* + 1)-mer *x_i–k:i_* is much smaller than the number of pseudo counts 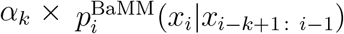, the higher order falls back to the lower order: 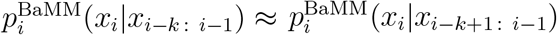. In this way, BaMMs adapt the order that is learned in a data- and motif position-specific fashion to the amount of data (*k*-mer counts) available. We assume that the correlation between nearby bases declines with their distance. This is reflected in the pseudo-parameters *α_k_* increasing with order *k*. For BaMMmotif2, we kept the same setting as in BaMMmotif, *α_k_* = 7 × 3^*k*^.

The motif model is optimized with the EM algorithm by maximizing the likelihood of the input sequences assuming zero or one motif occurrence per sequence (the ZOOPS model). It models the bound sequence using a *K*th-order inhomogeneous BaMM 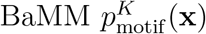 (where **x** = *x*_1:W_ is the binding site), and models the other unbound sequence regions using a *K′*th-order homogeneous BaMM 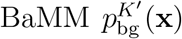 (*K′* is 2 by default). This background sequence model is trained by default on the input sequences. Potential binding site sequences **x** are ranked by their score 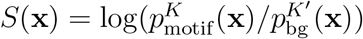. In the weak binding limit, this score is proportional to the Gibbs free energy Δ*G* of binding (Supplementary Section 1.5).

#### Learning positional binding preferences

BaMMmotif2 can learn the positional binding preferences for enriched motifs with respect to the center of the input sequences. By aligning the sequences around some anchor feature, such as a transcriptional start site, a 3’ splice site, or a binding site of some other transcription factor, the distance preference between enriched motifs and the reference feature can be learned. We parameterize the positional probability distribution with one parameter per position and ensure smoothness by adding *L_2_* penalties for the differences between successive sequence positions (Supplementary Section 1.5).

#### Masking sequences during the motif refinement stage

Sequences from *in vivo* experiments such as ChIP-seq commonly contain several distinct motifs from other TFs that together co-regulate their target genes. This can create two types of problems during the refinement stage. First, instead of refining the motif from the seeding stage, the model in some cases tends to learn two or even more motifs in the same higher-order model, as this often improves the likelihood on the training data. Second, if the seed motif is less enriched or less informative than other motifs in the positive sequence set, the model can switch from the seed motif to these other motifs. In this way, the weaker motif is not discovered at all. To avoid these two problems, we introduced a masking step in the EM optimization. We score all possible motif start positions in the input sequences using the PWM passed from the seeding stage to the refinement stage. We mask out all but the top *X*% of positions (X = 5 in this study) and ignore these positions in the EM iterations of the refinement stage.

### Motif assessment using average recall (AvRec)

To assess the performance of a classifier such as a motif model, one often plots the true positive predictions (TP) versus the false positive predictions (FP) over all score thresholds. Normalizing FPs and TPs to a maximum of 1 by plotting the true positive rate TPR = TP/Positives versus the false positive rate FPR = FP/Negatives yields the receiver operating curve (ROC). The often-used area under the ROC curve (AUC) is not a good quality measure for a motif model because in many applications the fraction of positive sequences (those carrying the motif) is much smaller than the number of negative sequences. When scanning the human genome for CTCF binding motifs in windows of 100 bp, for example, the ratio is about 1:30 (Fig. 6). At this ratio, a false discovery rate FDR = TP/(TP + FP) below 50% requires a ratio FPR/TPR < 1/30. So 29/30 = 97% of the ROC plot, the part with FDR > 50%, would be irrelevant. A predictor could have an AUC of 95% and never reach an FDR below 50%.

We therefore previously developed the Average Recall (AvRec) score [33], which averages the recall (the same as true positive rate and sensitivity) over a range of TP:FP ratios from 1:1, corresponding to FDR = 0.5, to 1:100, corresponding to FDR = 1/101 (Fig. 2 A). The AvRec score therefore considers the range of FDR most relevant in practice and has the additional benefit that a different positive-to-negative ratio than 1 simply results in a vertical shift of the AvRec curve on the logarithmic *y* axis.

**Figure 2.**
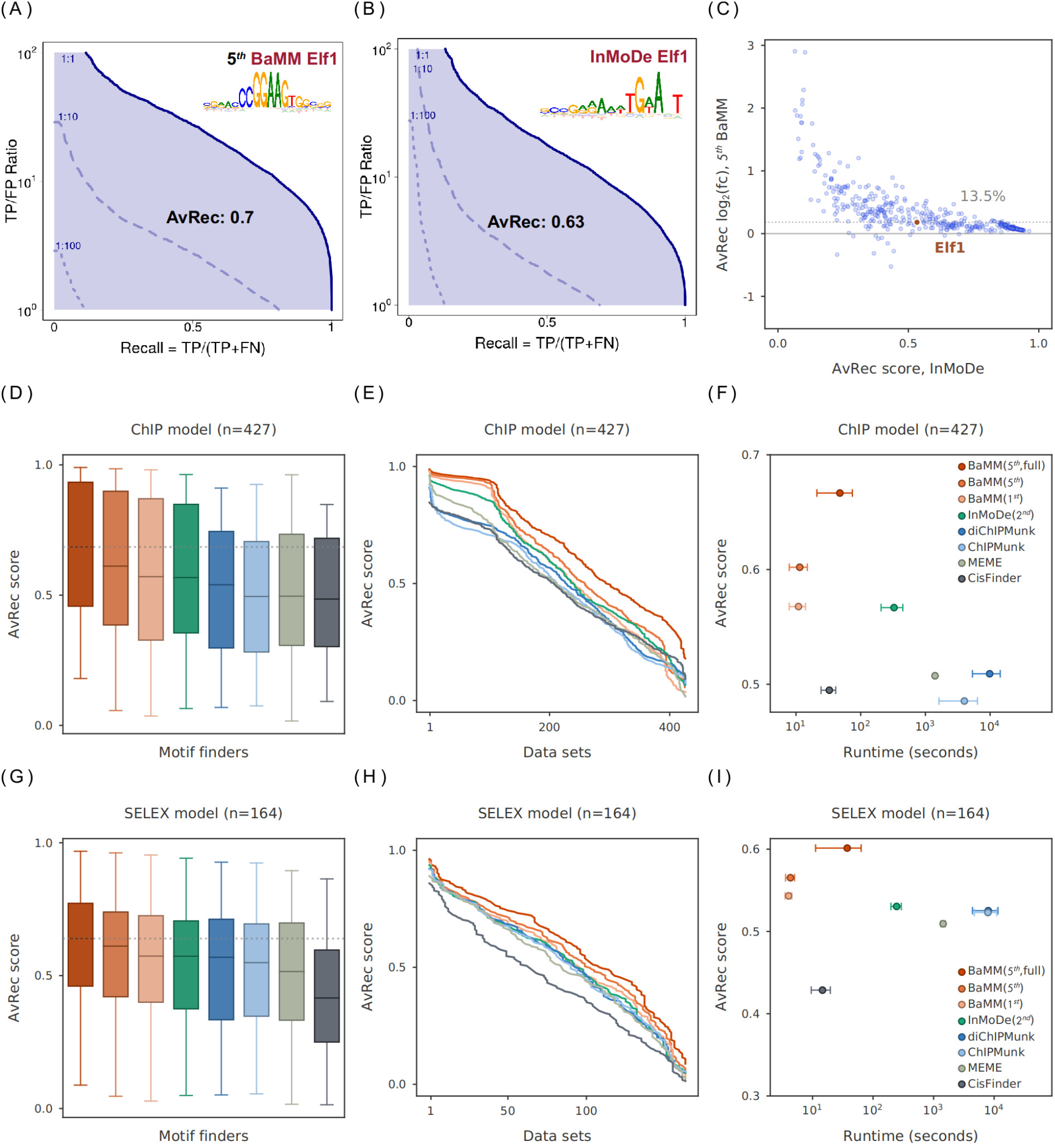
Performance of *de novo* motif discovery tools on *in vivo* and *in vitro* datasets. (**A**) AvRec analysis for fifth-order BaMM on the Elf1 ENCODE dataset. The AvRec is the recall averaged in log space over TP-to-FP ratios between 10^0^ and 10^2^. This ratio range corresponds to a precision between 1/(1 + 1) and 100/(1 + 100) = 0.99. Bold line: 1:1 ratio of positives to negatives. At 1:10 ratio (dashed) and 1:100 (dotted), the curves are shifted down by a factor of 10 and 100, respectively. Inset: motif logo of Elf1. (**B**) Same as (**A**) for the InMoDe model of Elf1. (**C**) log_2_ of AvRec fold change between fifth-order BaMMmotif2 and InMoDe models versus the AvRec of InMoDe. Each dot represents one dataset. Elf1 is highlighted in red. The median log2 fold change is 13.6%. (**D**) AvRec distributions as box plot, with boxes indicating 25%/75% quantiles and whiskers 95%/5% quantiles. Color code: see the legend in (**F**). (**E**) Cumulative distribution of AvRec scores on the 427 datasets. (**F**) Average runtime per dataset on four cores versus the median AvRec score. InMoDe and (di)ChIPMunk are not parallelized and ran on a single core. Whiskers: ±1 standard deviation. BaMM (5^th^, full): no masking step. (**G**-**I**) Analogous to (**D**-**F**) but for 164 HT-SELEX datasets from the Taipale lab [38].

To calculate the AvRec score, we first simulated 10-fold more negative than positive sequences using a second-order Markov background model learned on the positive set. We computed the motif scores 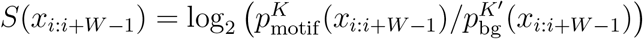 for all possible binding positions *i* (excluding the masked positions) and took the best score for each sequence. All sequences are sorted by descending score. The false positive count FP is the cumulative number of sequences from the negative set above the score cut-off, and TP is the cumulative number of positive sequences above the score cut-off.

### Benchmark design

The performance of BaMMmotif2 was evaluated together with five state-of-the-art motif discovery tools, MEME [32] as the most cited tool, CisFinder [34] for its speed and ability to run on large datasets, ChIPMunk [35] and diChIPMunk [27], which are used for generating the PWMs and dinucleotide PWMs in the HOCOMOCO database [36], and InMoDe [24], which can learn inhomogeneous Markov models of order 2 and beyond.

The processing of the ChIP-seq and HT-SELEX data is described in detail in Supplemental Material II. The motif discovery tools were run on the input sequence sets with default parameters, and four CPU cores were used for tools that could be parallelized (CisFinder, MEME, and BaMMmotif2). For tools that learn multiple motif models from one dataset, the motif models ranked top by the tools were evaluated in the 5-fold cross-validations and cross-cell line validations, while the best motif model out of maximal top 3 models for each dataset was evaluated in the cross-platform validations.

To assess the model performance over the given sequences, we first performed the benchmark on both human ChIP-seq [37] and HT-SELEX datasets [38] using 5-fold cross-validation (Fig. 1 I). The cross-cell-line validation was applied to ChIP-seq data (Fig. 1 II) and the cross-platform validations were applied to both ChIP-seq data HT-SELEX data (Fig. 1 III and IV). A more detailed description including tool settings can be found in Supplementary Section II.

## RESULTS

### Model performance on *in vivo* and *in vitro* data

We learned *de novo* motifs with each of the six tools on 427 ChIP-seq datasets for 93 transcription factors from the ENCODE project [37] and evaluated their performance using 5-fold cross-validation (Fig. 1I).

As an example, we compare in Fig. 2A and 2B the AvRec plot of a fifth-order BaMM with a second-order InMoDe model for the Elf2 motif, trained and tested on 5000 sequences of length 208 bp via 5-fold cross-validation. At a positives-to-negatives ratio of 1:1 (bold blue line) and a TP:FP-ratio of 10:1 (see *y* axis, corresponding to an FDR of 1/11), the BaMMmotif2 model achieves a recall of 0.81 and the InMoDe model achieves 0.69. At a positives-to-negatives ratio of 1:10 and a TP:FP-ratio of 10:1 (broken blue line), or, equivalently, at a positives-to-negatives ratio of 1:1 and a TP:FP-ratio of 100:1, the models achieve recalls of 0.12 and 0.13, respectively. When comparing AvRec scores between fifth-order BaMMs with second-order models from InMoDe across all 427 ChIP-seq datasets, BaMMs attain higher AvRec scores for 415 (97%) of the datasets, and the median AvRec of BaMMs is 13.6% higher than the one of InMoDe models (Fig. 2C).

Overall, the PWM-based tools, CisFinder, MEME, and ChIPMunk, are outperformed by the tools using higher-order models. BaMMmotif2 with first-order models performs on par with InMoDe and better than the first-order tools diChIPMunk. Fifth-order BaMMs achieve even better AvRec scores, as seen in the box plots and AvRec cumulative distributions of Fig. 2D and 2E, and in one-on-one comparisons in Fig. S2A. We also compared BaMMmotif2 with our previous tool BaMMmotif [28]. BaMMmotif2 is 10 times faster while being slightly more sensitive (Fig. S3).

Tools that learn higher-order Markov models can learn several motifs in one model, profiting from signals that are merely correlated with the real binding sites [29,39]. To find out whether BaMMs are affected or not, we introduced a masking step in the initial iteration of the EM algorithm (see Materials and Methods). We restrict the model refinement with the higher order BaMM to the 5 % potential motif positions with the highest scores scanned by the seeding PWM. In this way, we avoid overfitting and also speed up the refinement by a factor of 10. However, this robustness is paid by a loss in motif model performance (Fig. S4 and S5). The performance decrease could be caused in part by the limitation of being unable to select better sites during the refinement that were too different from the seeding motif, and in part because sometimes the BaMMs would otherwise have learned more than one distinct motif in a single model. To be on the conservative and robust side, we adopted the masking step in BaMMmotif2 for all our benchmarks in this study, unless explicitly stated otherwise.

Next, we assessed the performances of selected tools on 164 *in vitro* HT-SELEX datasets for 164 TFs [38]. Each dataset contains long oligomers of 200 bp. We also sampled 10-fold background sequences using the trimer frequencies from the same input set for estimating true negatives.

For the *in vitro* benchmark we observed overall similar trends as on the ChIP-seq data (Fig. 2G-I). CisFinder tends to learn longer motifs than the other tools, which probably helped it on the ChIP-seq data but hurt its performance on the HT-SELEX data. The BaMMs learned without masking (BaMM 5^th^, full; red) gained only 5% on the masked version (BaMM 5^th^; orange), whereas the gain had been 12% on the ChIP-seq data. This comparison shows that, on the ChIP-seq data, the fifth-order BaMMs trained without masking indeed tend to learn also motifs of co-occuring TFs that help to distinguish positive from negative sequences. If the goal is to learn the pure binding affinity of the ChIPped TF, masking should therefore be turned on for *in vivo* data.

### Assessing consistency of motif models across cell lines

ChIP-seq measurements have cell-type-specific biases associated with difference in chromatin accessibility, in particular of enhancers and promoters, and differences in TF concentrations [40]. A motif model that predicts only the binding affinity of the ChIPped TF should also perform well in predicting binding sites of the factor in other cell lines, whereas a motif model that has learned also motifs of co-occurring TFs and other sequence features with no direct effect on the binding affinity of the main TF should generalize badly to other cell lines in which different TFs will often co-occur with the ChIPped TF.

We therefore conducted a cross-cell line benchmark on *in vivo* data. We assessed the performance of models learned on ChIP-seq data from one cell line and tested on ChIP-seq data of the same TF from another cell line. We found 119 pairs of ChIP-seq datasets in the ENCODE database in which the same TF had been ChIPped in two different cell lines. We trained the model on one dataset and tested it on the other, and vice versa, resulting in 238 AvRec scores (Fig. 3).

**Figure 3.**
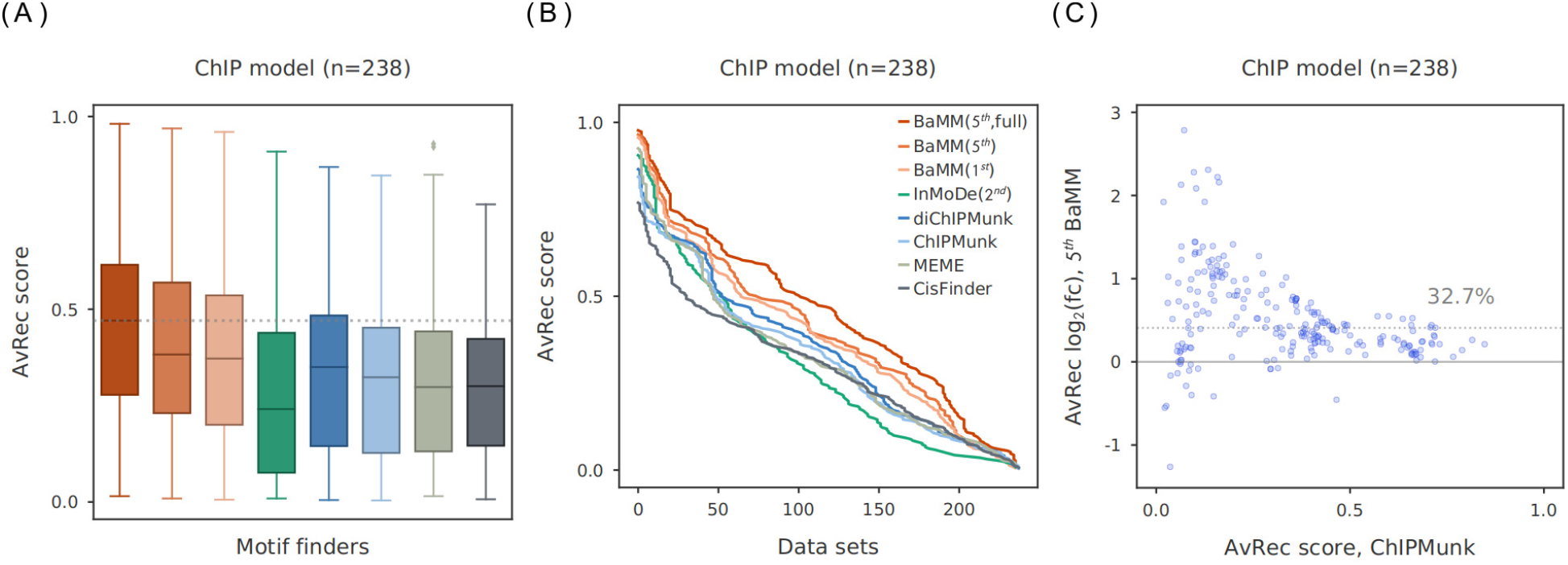
Cross-cell-line validation. 119 pairs of ENCODE datasets were used in this benchmark in which the same TF had been ChIPped in different cell lines. (**A**) AvRec distributions for 2 × 119 models that were tested on a ChIP-seq dataset from a different cell line than they were trained on. (**B**) Cumulative distributions of AvRec scores. (**C**) Log2 fold change in AvRec between fifth-order BaMMs and ChIPMunk for each of the 238 datasets. The median improvement is 32.7%.

Remarkably, the AvRec scores are around 0.2 lower for all tools than the AvRec scores in Fig. 2D obtained when training and testing in the same cell lines, with the PWM-based tools going from AvRec 0.5 to as low as 0.3. This quite dramatic decrease indicates that all models, even the simple PWMs, do not perform well for predicting bound sequences in another cell line. Remarkably, except for InMoDe, the predictive power of the higher-order models does not suffer more than that of the PWMs. This indicates that the higher-order models (except InMoDe) do not tend to overfit to sequence features that are specific to one cell line, such as co-occurring TFs. It is surprising that the fifth-order BaMMs trained without masking maintain or even improve their edge on the other models, despite our expectation that they would be the most prone to overfit on cell type-specific features.

### *In vitro* models predict *in vivo* binding and vice versa

Each measurement for detecting TF-DNA interactions has its own biases. ChIP-seq has biases from sequence-dependent PCR amplification, cell-type-specific sonication bias, and chromatin structure [41–43], while HT-SELEX has biased nucleotide compositions and depleted palindromes as a result of the library preparation, as well as sequence carry-over bias in selection cycles [31,44]. To assess how much models base their predictions on technical biases that would improve their performance when tested on the same platform but decrease their performance when tested on a different platform, we performed two cross-platform benchmarks.

First, for each of the tools we trained motif models on each of the 140 ChIP-seq datasets for which a HT-SELEX dataset with the same TF, but unnecessarily from the same cell line, was available. We discovered that several datasets were of too low quality to give reliable models. We therefore selected the 92 ChIP-seq datasets for which at least one of 8 tools achieved an AvRec score of ≥ 0.1 with the best model if multiple motif models are discovered. The first- and fifth-order BaMMs achieve better accuracies than the PWM-based models (Fig. 4A and 4B). ChIPMunk and diChIPMunk were at a disadvantage because they only predict one motif per dataset, while other tools may discover several motif candidates, the best out of the top three of which was evaluated. Fifth-order BaMMs trained without masking achieved better AvRec scores than those trained on masked ChIP-seq data, indicating that on ChIP-seq data masking might prevent training on weaker motif occurrences masked out by the less sensitive masking step. BaMMs also benefits from using higher than first orders on *in vivo* data.

**Figure 4.**
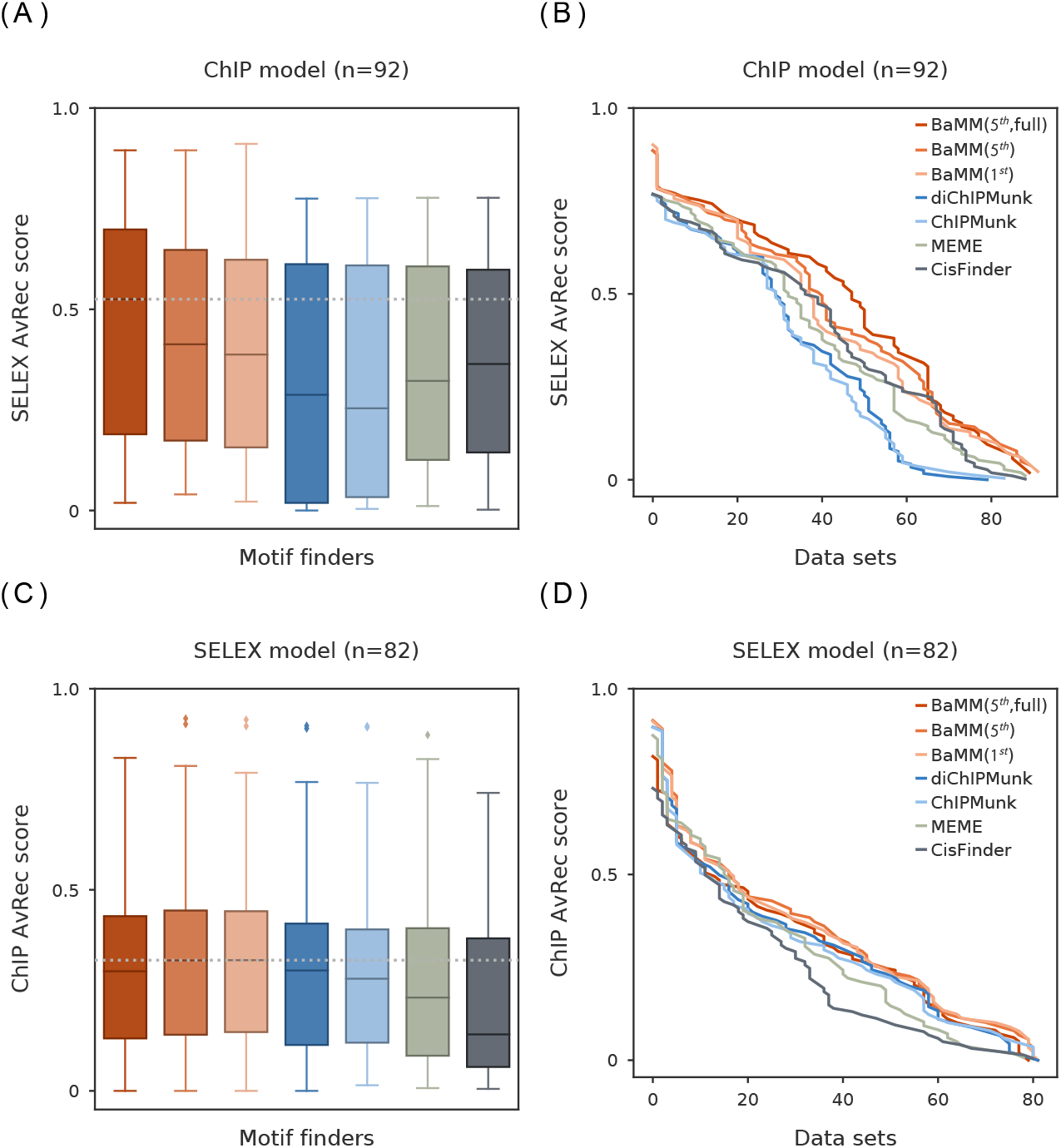
Cross-platform validation. (**A,B**) AvRec distributions and cumulative distributions for 92 models trained on ChIP-seq datasets and tested on HT-SELEX datasets for the same TFs using different tools. (**C,D**) Same as (A,B) for 82 motif models but trained on HT-SELEX datasets and tested on ChIP-seq datasets.

Second, for each of the tools we trained a motif model on each of the 82 HT-SELEX datasets for which a ChIP-seq dataset with the same TF was available. We selected the HT-SELEX datasets for which at least one of the 8 tools achieved an AvRec score of ≥ 0.1. Again, BaMMs achieved the best AvRec scores. However, we observed no major improvements from first to fifth order (Fig. 4C and 4D). This time, the improvements over PWM-based models are minor. When training fifth-order BaMMs on the HT-SELEX data, the masking step reduces model quality significantly, indicating that weak motifs not selected by the less sensitive masking stage are important to learn a good fifth-order model.

BaMMs learned similar information content in the first-order on ChIP-seq and on HT-SELEX data while showing no tendency to learn systematic biases of these platforms (Fig. S7). This demonstrates how the information in the first-order can help to improve cross-platform predictions.

### Extended flanking regions increase motif prediction accuracy

Various studies have shown that the flanking regions outside of the core binding sites affect TF binding, by affecting DNA shape preferences or by harboring binding sites of co-cooperatively binding TFs at variable spacings [14, 45–47]. Therefore, we investigated the impact of extending the core motifs, by adding two or four nucleotides on each side in the seeding motifs and refining the extended motifs with BaMMmotif2.

We find that for BaMMs trained on ChIP-seq datasets, extending the models by 2 × 2 or 2 × 4 positions indeed improves the motif model performance across all orders, and more so with increasing order (Fig. 5A). The improvement from no added positions to 2 × 4bp added is by 3% for zeroth order BaMMs (PWMs) and by 11% for fifth-order BaMMs (Fig. 5B and Fig. S8A). This indicates that flanking regions carry information mostly in the higher orders and not much in preferences for specific nucleotides.

**Figure 5.**
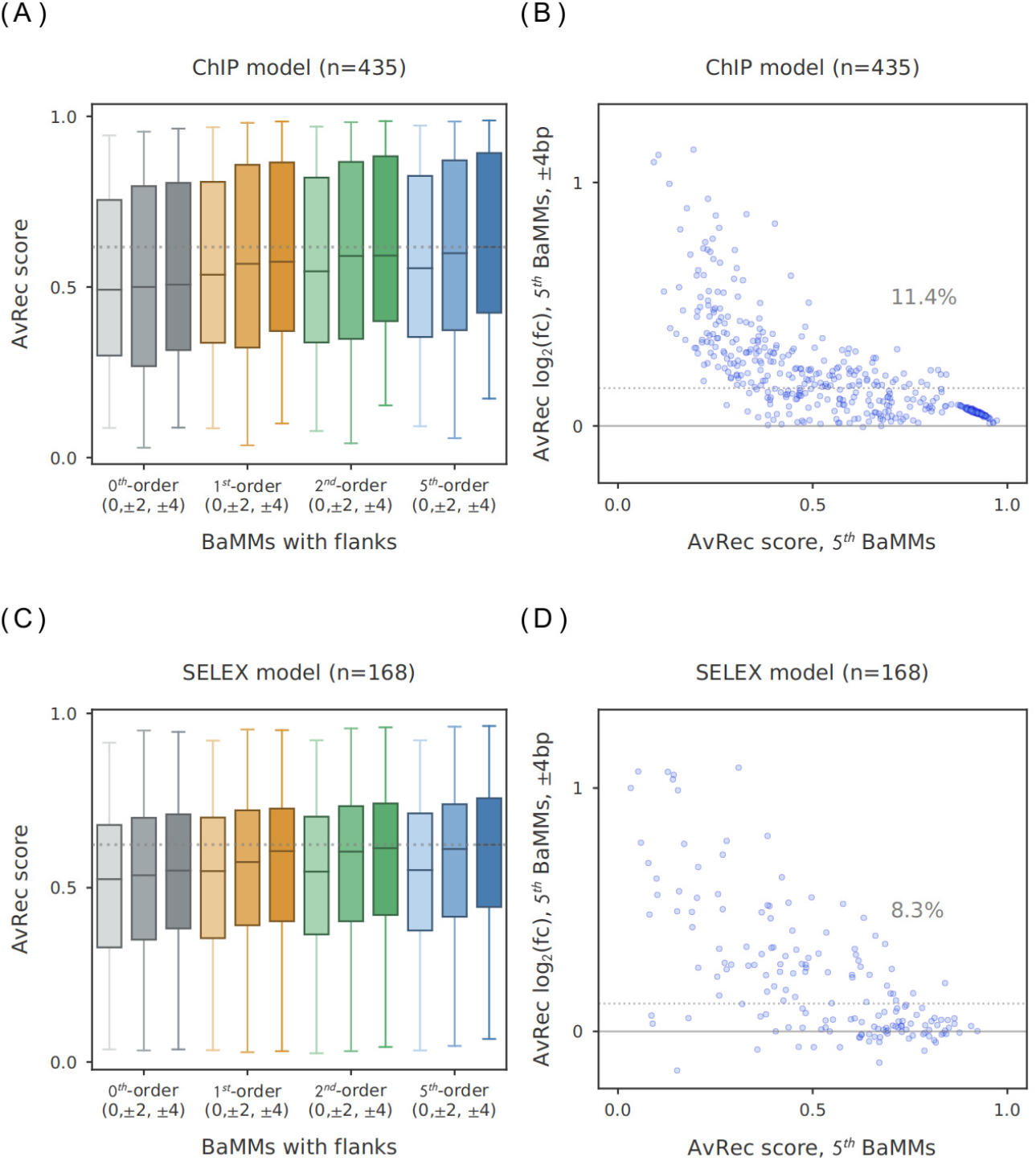
Extending the core motif by flanking positions improves motif performance. AvRec of BaMMs with different numbers of flanking positions added to the core motif, tested by 5-fold cross-validation. (**A**) AvRec distribution on 435 ChIP-seq datasets for models of order 0, 1, 2, and 5, each for three sizes of flanking regions: 0 bp, ± 2 bp, and ± 4 bp. (**B**) Log_2_ of fold change between fifth-order BaMMmotif2 models with ± 4 bp flanking positions and no added flanking positions. The median AvRec increase is 11.4 %. (**C,D**) Same as (A,B) for 168 HT-SELEX datasets.

It is not clear, however, if these improvements are due to DNA shape preferences that are reflected by preferences for certain di- and tri-nucleotides or by other sequence features of the genomic sequences such as motifs of co-occurring TFs. We therefore repeated the same analysis on HT-SELEX data. We restricted ourselves to long oligonucleotides of 200 bp because short oligonucleotides of 20 bp to 40 bp might not reflect well enough the physical properties of genomic DNA.

The results on the HT-SELEX data are very similar to those on ChIP-seq data (Fig. 5B and 5D). Again, PWMs gain much less AvRec score through 2 × 4bp extensions than fifth-order BaMMs (1.3% versus 8%, shown in Fig. S8B and Fig. 5D). This result confirms that the features picked up by the higher orders are not chiefly ones that are specific to genomic sequences but are also learned on in *vitro*-selected sequences and are therefore likely to be associated with DNA structural preferences.

### Learning positional binding preferences

Motifs often have certain positional preferences with regard to other motifs or genomic landmarks such as transcription start sites. Therefore, we introduced the possibility to learn the probability distribution of motif positions from the input data (Fig. S1A). Learning the positional distribution of motifs around ChIP-seq peak positions did not improve the median motif performances (Fig. S1B and S1C), probably because the information content of the positional distribution is very low when the the distribution is not much narrower than the window size. (The information content can be calculated as the difference between the entropies of the two positional distributions.) The positional preference is likely to have a positive impact when positioning effects are stronger, such as for splicing motifs around splice sites, core promoter motifs around transcription start sites, or TF binding sites of cooperatively binding TFs.

### Fifth-order BaMM predicts 1.8 million high-confidence human CTCF sites

The CCCTC binding factor CTCF plays a key role in transcriptional regulation through the formation of chromatin loops and topologically associated domains (TADs). Chromatin loops and TADs are believed to be formed by DNA getting extruded through a cohesin ring [48]. The extrusion and loop growth are halted when the cohesin encounters CTCF bound to each of the two strands of the DNA in the preferred orientation [49].

Zhang et al. [50] identified 112 thousand CTCF binding sites from the GM12878 cell line, using a PWM for CTCF motif from the JASPAR database (ID: MA0139.1) [51]. We took the ENCODE dataset for CTCF from the same cell line, trained a fifth-order motif with the length of 67 bp in total, a core of 27 bp with 2 × 20 bp flanks (Fig. 6A) and scanned the human genome (hg19) with it using the script BaMMScan in the BaMMmotif2 software. Fig. 6B shows a histogram of the BaMMmotif2 score distribution obtained on the human genome (orange).

**Figure 6.**
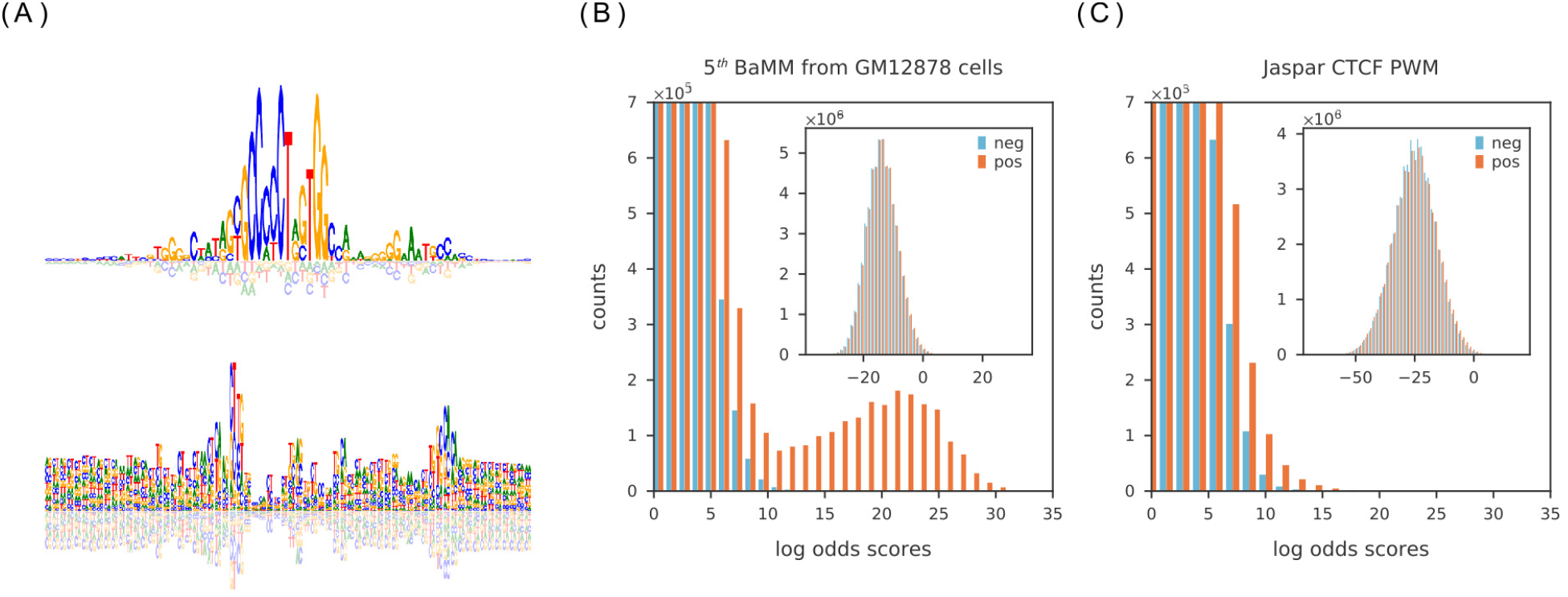
A fifth-order BaMM predicts 1.8 million high-confidence CTCF sites in the human genome. (**A**) Information content of CTCF motif model in zeroth- (top) and first-order (bottom). Distributions of motif log-odds scores on the hg19 genome (orange) and the reverse genome (blue), scanned with a fifth-order BaMM learned from the GM12878 cells (**B**) and a PWM (JASPAR ID: MA0139.1) (**C**). BaMMmotif2 predicts 1.8M CTCF binding sites with log odds scores > 11 at a FDR below 0.5% (B), the JASPAR PWM predicts 67k binding sites at a FDR of 15% (C).Inset: the full distributions of the predicted CTCF sites on the whole genome (orange) and the reverse genome (blue).

Remarkably, the motif score distribution on the human genome displays a two-peaked distributions, with a minimum around score 11 separating a small peak from the bulk of the distribution. This shape is suggestive of a mixture of two somewhat overlapping but separated components, a huge component comprising the non-CTCF sites and a small component with true CTCF sites.

If this is true, we would expect few of the sites with scores above 11 to be false positive predictions. To verify this, we estimated the number of false positive predictions per score bin by scanning the reversed human genome sequence with the CTCF BaMM. The score distribution is shown with blue bars in Fig. 6B. The distribution of negatives show only 7542 positions with scores above 11, while the distribution of scores on the human genome has 1.8 million positions. This confirms our hypothesis that these 1.8 million positions in the human genome are high-confidence CTCF sites with a FDR of 7542/(1.8 × 10^6^) < 0.5%.

To compare these predictions with a state-of-the-art prediction using the CTCF PWM in the JASPAR database (ID: MA0139.1) [51], we have plotted in Fig. 6C the scores distributions of the PWM (obtained with BaMMscan with second-order background Markov model) on the human genome (orange) and the reversed human genome. While the negatives look very similar to the one for the BaMM, the score distribution on the human genome has no separate peak at high scores, and only 67 thousand positions have a score larger than 11 (FDR < 0.15).

These results indicate that higher-order BaMMs can predict many more binding sites for CTCF and other factors with high confidence than PWM-based methods.

## Discussion

We presented BaMMmotif2, a fast and accurate *de novo* motif discovery algorithm for large-scale transcriptomic data. BaMMmotif2 builds on our earlier theory of Bayesian Markov models (BaMMs) implemented in BaMMmotif. BaMMs employ pseudocounts from model order *k* − 1 to stabilize the estimation of the conditional probabilities for order *k*, for all orders *k* from 1 to the maximum order (five in this study). In this way, they can learn higher orders if a sufficient number of *k*-mer counts was observed to estimate them but otherwise fall back to a lower order that can still be estimated safely.

BaMMmotif2 was written from scratch in C++ using explicit AVX2 vectorization and multicore parallelization. We developed a novel, fast seeding method to find enriched patterns that scales almost independently of the input set size. We also added a masking step to force the refinement stage to only refine the seed motifs and prevent it from learning in addition other predictive features such as co-occurring motifs of other TFs or experimental sequence biases. We also developed a Bayesian approach to learn position binding preferences from the input data.

By their sheer number, ChIP-seq datasets are the dominant source of information for TF binding affinities. Therefore, most benchmark comparisons of *de novo* motif discovery tools have been performed exclusively or predominantly on ChIP-seq data. However, for assessing the quality of models more complex and informative than PWMs, such as higher-order Markov models and mixture models, ChIP-seq data are problematic for several reasons. First, they often have complex sequence biases [40], which higher-order models can learn to distinguish from negative sequences generated with random background models. To alleviate this problem, second order background models should be used, but even this might be insufficient to eliminate learning generic sequence biases of the ChIPped versus random sequences. Second, sequences in ChIP-seq peaks usually contain in addition to the motif of the ChIPped TF the binding motifs of co-binding factors [39]. Complex models can improve their predictive performance by scoring sequences highly that contain any of these co-occuring motifs. This is possible even within a short motif length by learning the motifs superposed with each other, with the higher orders preventing mixing and blurring of motifs [29]. Although improving the apparent model performance, such models do not describe faithfully the binding affinity of the ChIPped factor.

Our goal was to compare PWM-based motif discovery tools with tools employing more complex models: dinucleotide weight matrices, parsimonious context trees, and BaMMs. We therefore set up a cross-cell line benchmark to assess how well the motif models learned in one cell line can predict binding in another cell line. Furthermore, we conducted a cross-platform benchmark, in which we trained the models on ChIP-seq data and tested them on HT-SELEX data, and vice versa. The results show that among the tested tools, those with more complex models still tend to perform better in these benchmarks, albeit with smaller improvements over the PWM-based tools. The improvements from higher orders were particularly marked for the BaMMs. So, most of the information in higher orders seems to be transferable between cell lines and measurement platforms.

Even though we did not see clear signs of overfitting in our BaMMs, we introduced sequence masking as a precaution against overfitting to other motifs and technology- or cell line-dependent sequence biases. We use the seed PWM to mask out all but the top-scoring 5% of positions, and we train the higher-order BaMM only on the remaining 5%. We thereby ensure that only sequence regions that actually carry the seed motif can be learned by the BaMM. The performance drop between training fifth-order BaMMs with and without masking was 8% on HT-SELEX data and 12% on ChIP-seq data (Fig. 2D,G, Fig. S4). This indicates that if higher-order BaMMs profit from learning co-occurring motifs at all, the effect on their performance is quite limited. Still, if the goal is to learn binding affinities and not just predict motifs from *in-vivo* sequence data, we recommend to run BaMMmotif2 with the masking. On *in-vitro* data, however, masking is not necessary and in order to make use of the 5% improvement we recommend to run BaMMmotif2 without masking. However, even with masking the fifth-order BaMMs still perform competitively with the state-of-the-art tools while being significantly faster.

Transcription factors combine base-with DNA shape readout [13]. Instead of studying the TF-DNA binding using only the sequence features, some models utilize DNA shape features predicted from the sequence to enhance motif models [52–54]. The shape descriptors these tools use, like minor groove width, helical tilt and bent, or propeller tilt, are predicted from five-mer tables computed using molecular dynamics calculations. Given enough data, it is therefore evident that higher-order models such as BaMMs can learn these DNA structural preferences implicitly, yet are not limited to the pre-defined shape descriptors.

In recent years, deep learning approaches have become popular for learning motif models with very good predictive performance [52, 55, 56]. Such models usually take advantage of contextual information such as co-occurring motifs, which increases their predictive power but serves a different purpose than the models we discuss here: learning a model for the sequence dependence of the binding affinity of a factor. In addition, BaMMs have the advantage or being conceptionally simple and interpretable in terms of *k*-mer dependent energy terms.

In conclusion, we have shown that higher order models for binding motifs improved binding site predictions on a large collection of ChIP-seq and HT-SELEX datasets, both in crossvalidated setting and when training and testing on different experimental platforms and cell lines. Importantly, clear improvements in predictive performance are even seen beyond first order models: BaMMs of fifth order show a solidly improved performance across the bench over the tested state of the art tools, while being significantly faster.

## Availability

### Data

#### ENCODE database

We evaluated the performance of selected algorithms on human ChIP-seq datasets from [the ENCODE portal](https://www.encodeproject.org/) [37] until March 2020. In total, there are 435 datasets for 93 distinct transcription factors. The top 5000 peak regions sorted by their signal value are selected for each dataset when peaks are more than 5000, and all peaks are chosen if there are fewer than 5000 peaks. Positive sequences are extracted ±104 bp around the peak summits. Background sequences are sampled by the trimer frequencies from positive sequences, with the same lengths as positive sequences and 10 times the amount of positive sequences. 8 datasets are excluded from all the results because diChIPMunk fails to learn models within 3 hours.

#### HT-SELEX datasets

For HT-SELEX data, we downloaded 164 datasets with 200 bp-long oligomers from Zhu et al. [38], which are deposited in the European Nucleotide Archive (ENA) under the accession PRJEB22684. Each dataset represents one non-redundant transcription factor. For each dataset, we selected 5000 sequences from each selection round without any sorting.

The HT-SELEX data contain reads from at least four selection cycles, and the measured binding affinity iteratively increases with the cycles. Thus, we chose the sequences from the fourth selection rounds with detected high affinities for motif training and testing in the main paper. Since ChIPMunk and diChIPMunk took longer than 2 hours to run on the full datasets, we selected 5000 sequences out of the millions of reads as training and test sequences. To examine the power of BaMMs in learning the weak binding sites, we also used sequences from the second and third selection rounds. Background sequences are sampled in the same way as described previously.

### Software and parameters

The new version of BaMMmotif2 software is implemented in C++ and Python3. The code is licensed under GPLv3 and freely accessible without registration at github.com/soedinglab/PEnG-motif, and github.com/soedinglab/BaMMmotif2, and supported on Linux and MacOS. They are also integrated into our [webserver](https://bammmotif.soedinglab.org/) [33].

### Results and analysis scripts

The analysis scripts are available in Jupyter Notebook format at github.com/soedinglab/bamm-benchmark.

## Acknowledgements

We thank Matthias Siebert and Anja Kiesel for help with designing the code structure of BaMMmotif2, the members of the Söding lab, especially Eli Levy Karin, Saikat Banerjee, Milot Mirdita and Ruoshi Zhang for discussions, and the genomics research community for sharing their data.

## Author Contribution

W.G. developed the BaMMmotif2 software. M.M., C.R. and J.S. developed the seeding stage software (PEnG). W.G. implemented the statistical approach and conducted all the benchmarks. W.G. and J.S. wrote the manuscript. J.S. supervised the research.

