## Supplementary Information for "Bayesian Markov models improve the prediction of binding motifs beyond first order"

### Part I

### Supplemental Figures

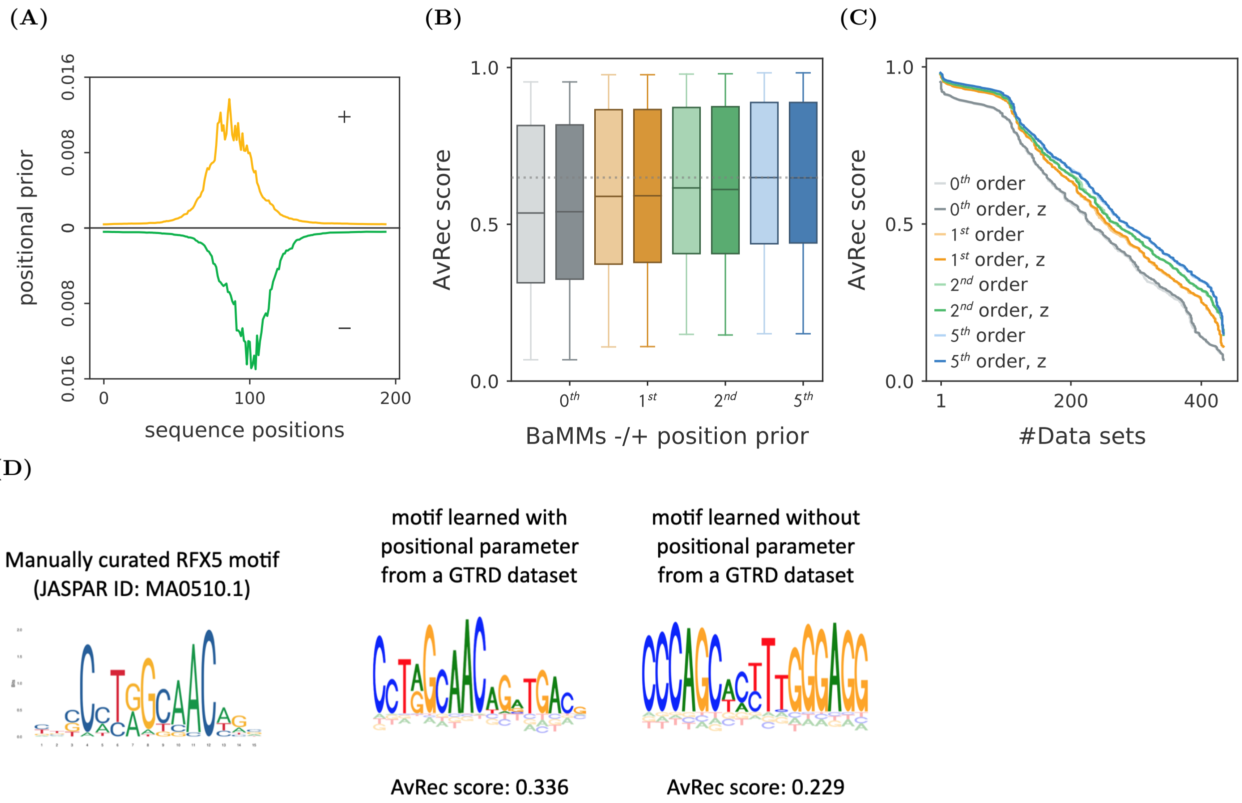


**Figure S 1. Optimization of positional prior on *in vivo* data.** BaMMs with zeroth- (grey), first- (orange), second- (green) and fifth-orders (blue) are trained and tested on 435 ENCODE datasets using 5-fold cross-validation. Panel (**A**) shows the distribution of optimized positional priors over the positions on both sequence strands that are center around ChIP-seq summits for GABP$\alpha$ motif. Panel (**B**) shows the AvRec distributions as box plot, with boxes indicating 25%/75% quantiles and whiskers 5%/95% quantiles. The colors are for different orders with- (dark colors) and without (light colors) optimization of positional prior in the motif training. Panel (**C**) shows the cumulative distributions of AvRec scores on 435 datasets. There is no major difference before and after the positional prior optimization. Panel (**D**) shows the motif for RFX5 factor learned from a GTRD dataset [1] with (middle) and without (right) the optimized positional parameter, compared to the reference motif reported in the JASPAR database [2] (left).

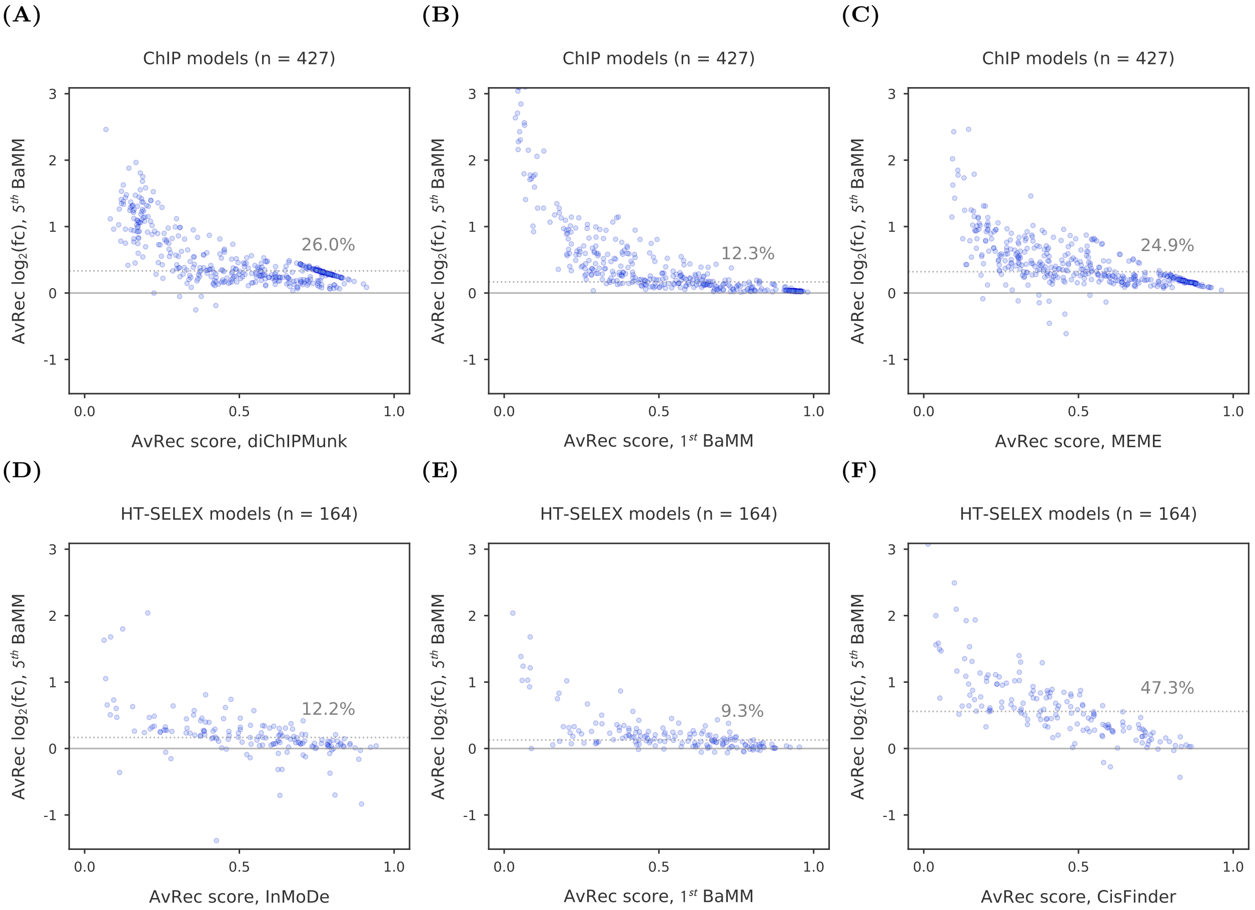


**Figure S 2. Performance comparison of motif discovery tools on *in vivo* and *in vitro* data.** $\log_2$ of fold change in AvRec between fifth-order BaMMmotif2 and diChIPMunk models versus AvRec of diChIPMunk models (**A**), first-order BaMMmotif2 (**B**) and MEME PWMs (**C**) on 427 ChIP-seq datasets. Each dot represents the test on one dataset from either ChIP-seq or HT-SELEX. The grey dashed lines indicate the median $\log_2$ fold change is 26%, 12.4% and 24.9% respectively. (**D-F**) Similar comparisons as (**A-C**) but on 164 HT-SELEX datasets.

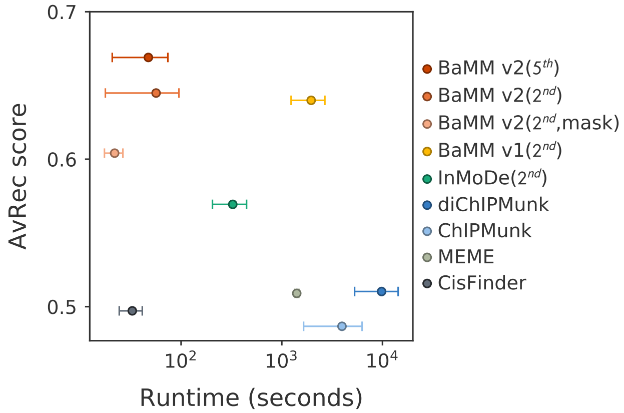


**Figure S 3. Benchmark on *in vivo* data.** Average runtime per dataset on a server with 4 cores versus the median AvRec score of several *de novo* motif discovery tools, including the previous version of BaMMmotif, validated on 419 datasets with 5-fold cross-validation, with MEME, CisFinder, BaMMmotif and BaMMmotif2 running on 4 CPU cores. Whiskers indicate the standard deviation of AvRec score. $2^{nd}$-order models trained using BaMMmotif and BaMMmotif2 have similar average AvRec scores, yet BaMMmotif2 is $>\tilde{1}0$ times faster than BaMMmotif.

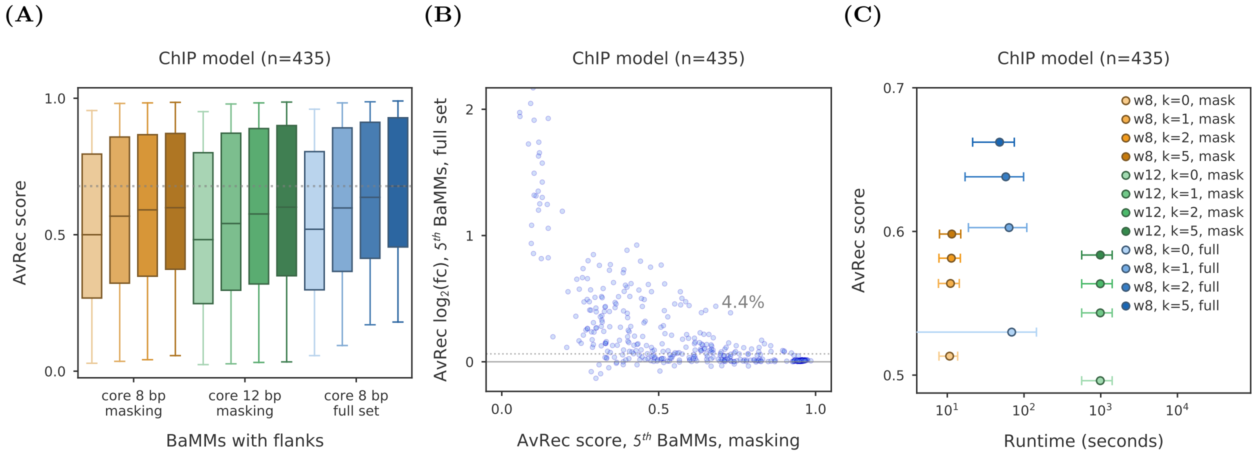


**Figure S 4. EM optimization using the full set compared to masking 95% sequences on ChIP-seq datasets.** (**A**) Using the full set of sequences for the EM optimization (blue) improves the performance of higher-order models, while extending the core regions for searching the enriched patterns (green) does not contribute to motif discovery, in comparison to that with 8 bp for seeding and masking 95% sequences for the EM optimization (yellow). All box-plot whiskers show 95th/5th percentile. Each cluster contains models with different orders (zeroth-, first-, second- and fifth-order). (**B**) Fifth-order BaMMs optimized on the full sequences set have a 4.4% AvRec fold increase compared to those trained only 5% sequences. (**C**) Using a masking step improves the speed by 10-fold, in comparison to using the full set for learning motif model.

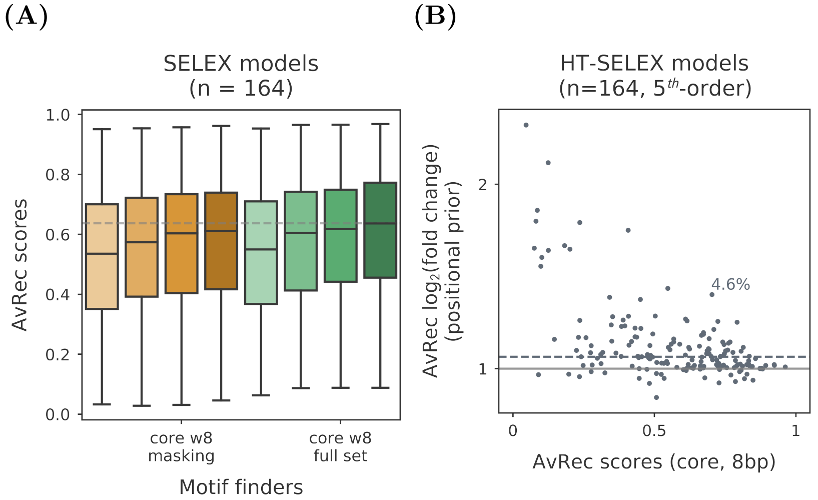


**Figure S 5. EM optimization using the full set compared to masking 95% sequences from HT-SELEX datasets.** (**A**) Using the full set for motif refinement (green) improves the performance of higher-order models over that using only 5% sequences (yellow). All box-plot whiskers show 95th/5th percentile. Each cluster contains models with different orders (zeroth-, first-, second- and fifth-order). (**B**) Fifth-order BaMMs with full set of sequences for optimization has a 4.6% AvRec fold increase compared to it with only 5% sequences for optimization.

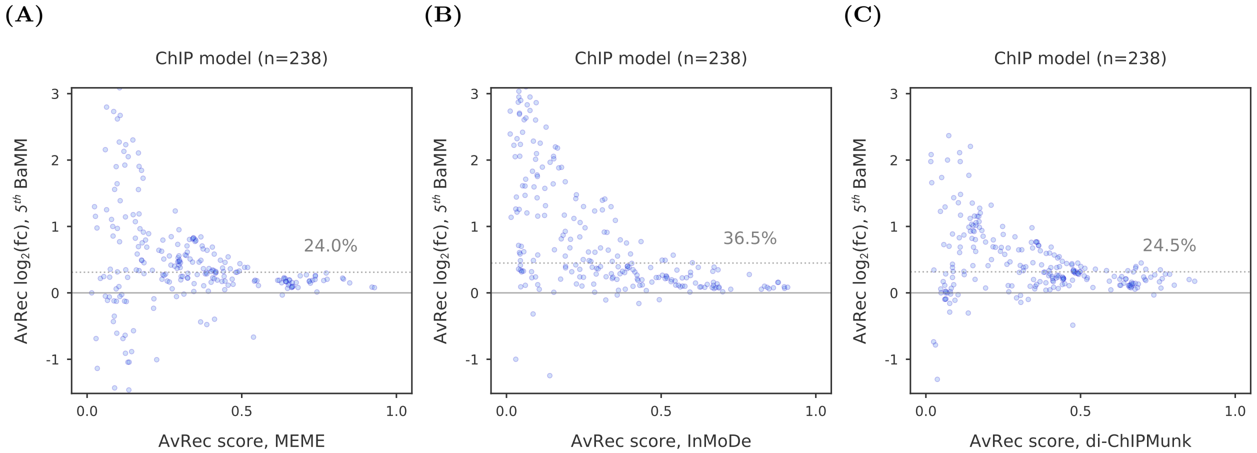


**Figure S 6. Cross-cell-line validation.** $\log_2$ of fold change in AvRec between fifth-order BaMMmotif2 and PWMs from MEME (**A**), second-order models from InMoDe (**B**), and first-order models from diChIPMunk (**C**), when comparing to AvRec scores of the latter models in 238 paired ENCODE datasets. Each dot represents one test. The range of AvRec scores is chosen from 0.5 to 8 and the outliers are not shown in these plots.

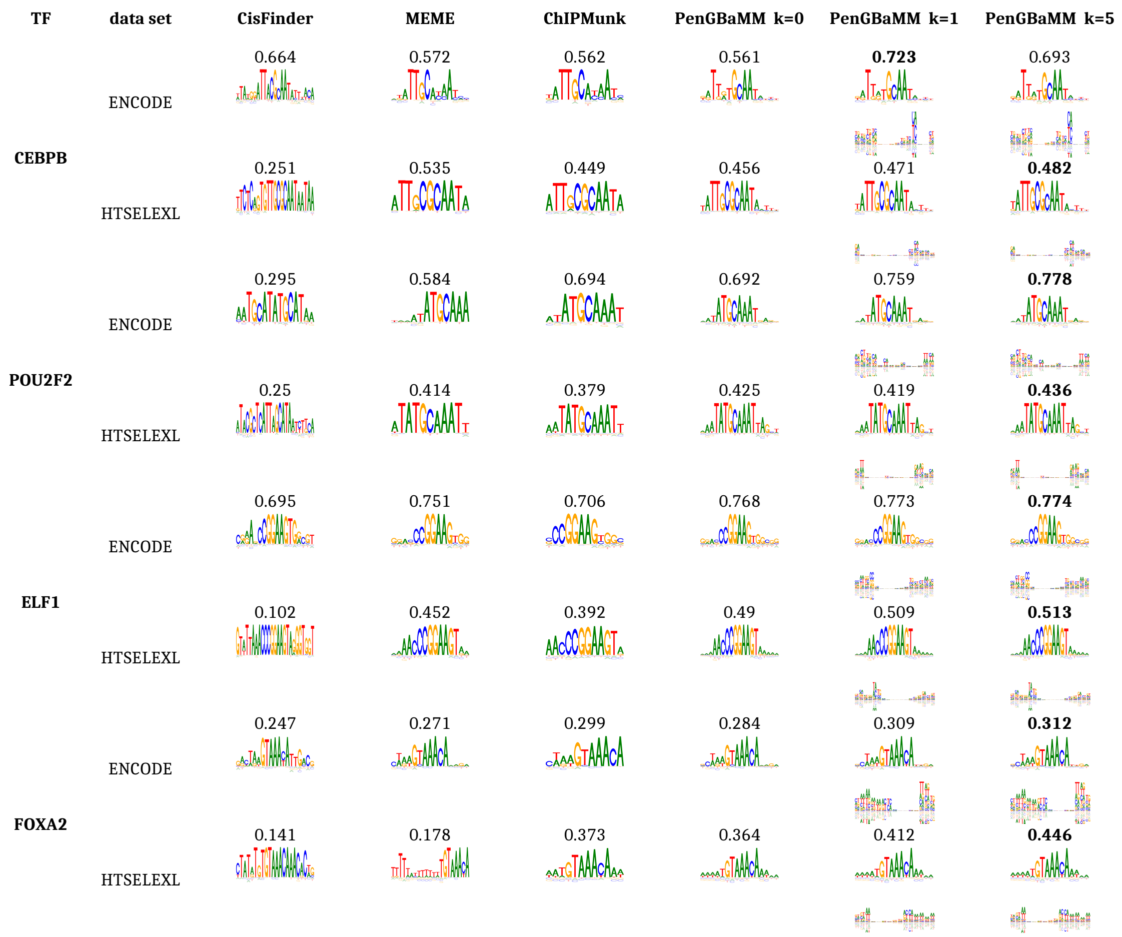


**Figure S 7. Sequence logos and AvRec scores of motifs models from cross-platform validation.** Motif models are trained by different models for four transcription factors: CEBPB, POU2F2, ELF1, and FOXA2. For each transcription factor, the first row shows models learned on ENCODE data by applying different tools. The number above each logo represents the AvRec score when testing the model on the corresponding HT-SELEX data. The second row shows the models learned on HT-SELEX data and AvRec scores when testing models on ENCODE data. For BaMM models, both zeroth- and first-order logos are plotted.

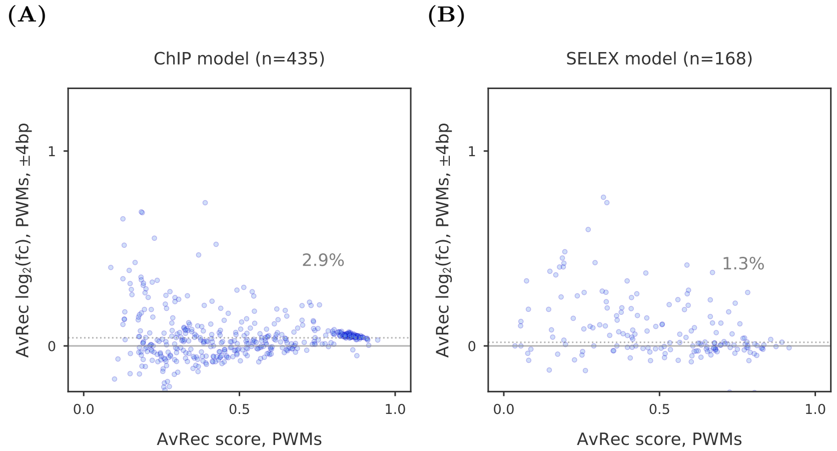


**Figure S 8. Impact of extending core motif regions on PWMs.** (**A**) $\text{Log}_2$ of fold change between PWM models with ± 4 bp flanking positions and no added flanking positions, using 435 datasets. Median AvRec change is 2.9 %. (**B**) Same as (**A**) but on 168 HT-SELEX datasets.

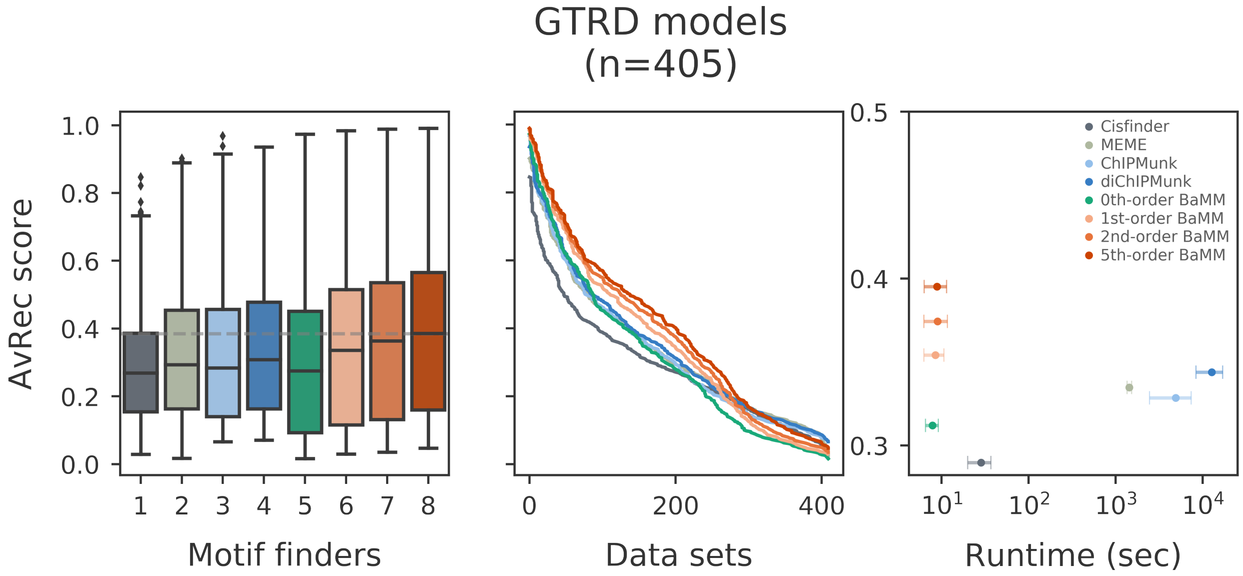


**Figure S 9. Quantitative performance on *in vivo* GTRD datasets.** The selected tools are applied to 405 GTRD datasets [1] and their Average Recall were calculated by 5-fold cross-validation, similar to **??**. (**A**) AvRec distributions as box plot. All box-plot whiskers show 95th/5th percentile. (**B**) The cumulative of AvRec scores on 405 datasets. (**C**) Average runtime per dataset on a server with 4 cores versus the median AvRec score. Whiskers: standard deviation.

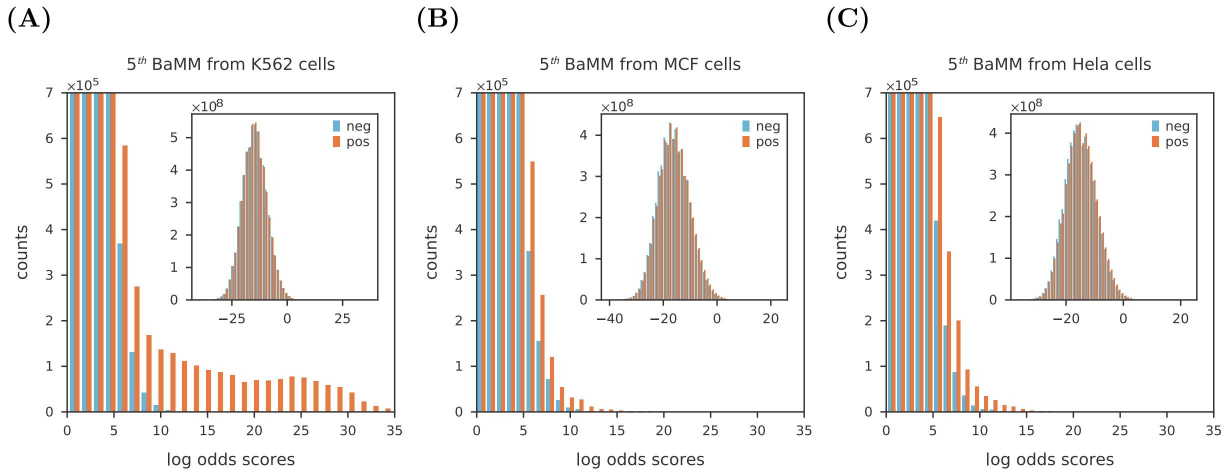


**Figure S 10. Distributions of CTCF motif scores across the whole human genome (hg19), scanned by different BaMMs** The distribution of log odds scores that were calculated on all the binding sites with a 5th-order BaMM of length 67 bp trained on ChIP-seq datasets either on K562 (**A**), MCF (**B**) or HeLa (**C**) cell lines on the whole genome (orange) and reverse genome (blue). The background model is a second-order Markov model. Inset: the full distributions of on the whole genome (orange) and reverse genome (blue). Note that the scores are the only 1% of the total that are randomly drawn from the full distribution of the whole genome.

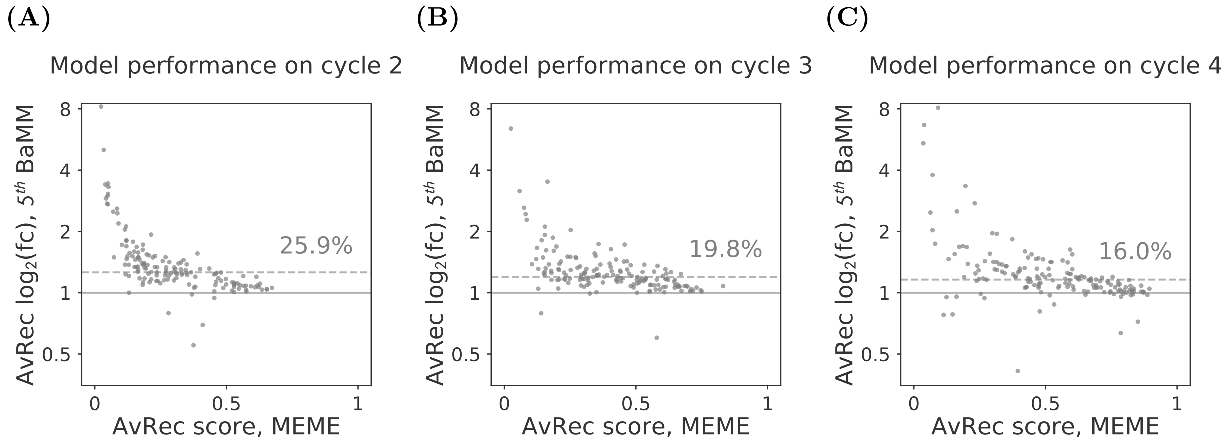


**Figure S 11. Performance comparison of BaMMmotif versus MEME on weak binding prediction.** $\log_2$ fold change between fifth-order BaMMmotif2 models and MEME models versus AvRec of MEME, with AvRec analyzed by 5-fold cross-validation on sequences from the 2nd- (**A**), 3rd- (**B**) or 4th- (**C**) selection cycle of 164 HT-SELEX datasets. The median fold change increases are 25.9%, 19.8% and 16%, respectively (grey dashed lines). Each dot represents one data set.

### Part II

### Supplemental Methods

The supplemental material provides further details of the theoretical basis, the implementation of the BaMMmotif2 package, and the processing procedure of the datasets that are used for this benchmark. It also documents the parameters used for testing the motif discovery tools in the benchmark. It ensures the reproducibility of the results in this paper.

#### 1 *De novo* motif discovery and refinement

##### 1.1 The fast seeding phase: PEnGmotif

We describe PEnGmotif (Pattern-based discovery of enriched genomic or transcriptomic sequence motifs), an efficient method for discovering sequence patterns enriched in a set of nucleotide sequences over random expectation sampled from a second-order background model. The enriched patterns found by PEnGmotif are optimized to PWMs and serve as seeds to initialise the refinement stage by BaMMmotif2.

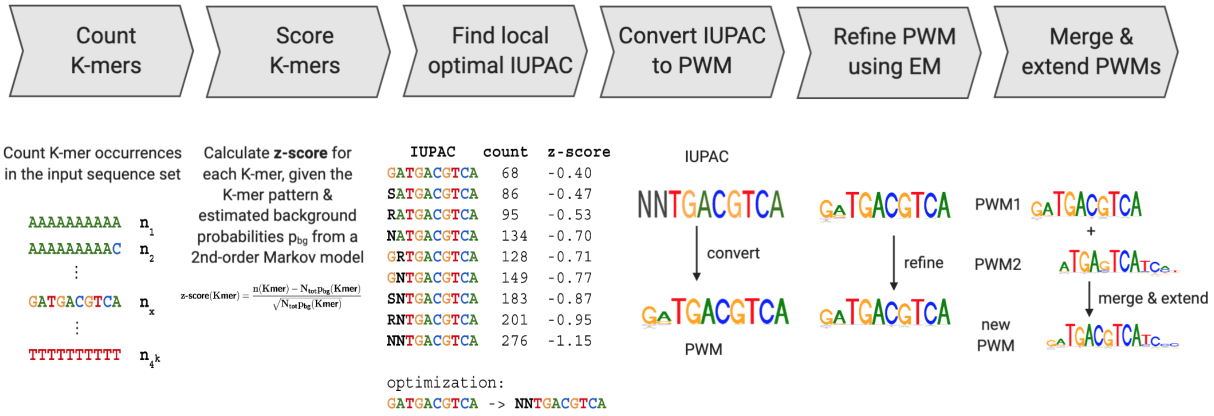


**Figure S 12. Workflow of the fast seeding stage.** Sequences from high-throughput assays, such as ChIP-seq, SELEX and PBM, are provided as input data. (i) Occurrences of all $K$-mers of a fixed specified length (default 10) are counted. (ii) An enrichment $z$-score is calculated for each $K$-mer based on a Poisson model. (iii) High-scored $K$-mers are optimized from the nucleotide alphabet (ACGT) to a degenerate IUPAC alphabet with 11 letters (ACGTRYSWSKN). (iv) The locally optimal IUPAC patterns are converted to PWMs. (v) PWMs are refined using the Expectation Maximisation algorithm. (vi) PWMs with similar overlapping regions are merged and extended.

Let $K$ be the length of patterns that will be analysed (e.g. $K = 8$ used in the study). First, the number of occurrences of each of the $4^K$ non-degenerate seed patterns of length $K$ are counted in a $4^K$-dimensional array with $\mathbf{x} \in \{\mathrm{A}, \mathrm{C}, \mathrm{G}, \mathrm{T}\}^K$. $p_{\mathrm{bg}}(\mathbf{x})$ denotes the probability of observing $K$-mer $\mathbf{x}$ in absence of specific binding. $p_{\mathrm{bg}}(\mathbf{x})$ can be directly counted from large background sequence sets or modelled as a homogeneous Markov model on a background data set or the dataset itself. For example, $p_{\mathrm{bg}}(\mathbf{x})$ is learned from the genomic input, a mock immunoprecipitation or the input sequence library prior to the selection in HT-SELEX. We model the background probability using a homogeneous Markov model of order $K'$ ($K' = 2$ by default):

$$p_{\text{bg}}(x_{i_0\,:\,i_1}) = \prod_{i=i_0}^{i_1} p_{\text{bg}}(x_i | x_{i-K'\,:\,i-1}). \tag{1}$$

We assume the number of occurrences in absence of specific binding to follow a Poisson distribution: $\mu = L_{\text{tot}} p_{\text{bg}}(\mathbf{y})$, where $L_{\text{tot}} = \sum_{n=1}^{N} (L_n - K + 1)$ is the total number of all counted patterns in the input sequences ($N$ is the total sequence number and $L_n$ is the length of $n'$th sequence).

**$z$-score** We compute $Z$-scores for all non-degenerate $K$-mer patterns. The $Z$-score is the deviation from expectation divided by the standard deviation. As for the Poisson distribution the variance equals the mean, the $Z$-score is:

$$Z(\mathbf{y}) = \frac{n(\mathbf{y}) - L_{\text{tot}} p_{\text{bg}}(\mathbf{y})}{\sqrt{L_{\text{tot}} p_{\text{bg}}(\mathbf{y})}}\,. \tag{2}$$

The z-score can be used to pre-filter what $K$-mers should enter the optimization routine.

**p-value** As we are also interested in highly enriched sequences (x) and (y) are fulfilled and we can use the Stirling approximation to calculate the p-value:

$$\begin{aligned}
\text{p-value}(\mathbf{y}) &= \sum_{k=n}^{\infty} \frac{\mu^k}{k!} e^{-\mu} \\
&= \frac{\mu^n}{n!} e^{-\mu} \sum_{k=0}^{\infty} \frac{\mu^k}{(n+1)\cdots(n+k)} \\
&\lessapprox \frac{\mu^n}{n!} e^{-\mu} \sum_{k=0}^{\infty} \frac{\mu^k}{(n+1)^k} \\
&\approx \frac{\mu^{n(\mathbf{y})}}{n(\mathbf{y})!} e^{-\mu} \frac{1}{1 - \mu/(n(\mathbf{y})+1)} \\
\log \text{p-value}(\mathbf{y}) &\approx n(\mathbf{y}) \log \frac{\mu}{n(\mathbf{y})} + n(\mathbf{y}) - \mu - \frac{1}{2}\log(2\pi n(\mathbf{y})) - \log\left(1 - \frac{\mu}{n(\mathbf{y})+1}\right).
\end{aligned} \tag{3}$$

**Mutual information** We optimize the mutual information (MI) between two random variables,

$$\text{MI}(q) = -qH(p_{\text{obs}}) - (1-q)H(p_{\text{exp}}) + H(p), \tag{4}$$

with $H(x) := -x\log x - (1-x)\log(1-x)$.

We then find locally optimal non-degenerate patterns with a recursive function which takes a $K$-mer $\mathbf{y}$ and checks for all its neighbouring $K$-mers, i.e. those that are at most one substitution away. If it finds a neighbouring $\mathbf{y}_{\text{neigh}}$ with a better mutual information, the function is called recursively with $\mathbf{y}_{\text{neigh}}$ as an argument. If no neighbour of $\mathbf{y}$ has better mutual information than $\mathbf{y}$, $\mathbf{y}$ is appended to the list of locally optimal $K$-mers. Similarly, we optimized the high-scored $K$-mers from the nucleotide alphabet (ACGT) to a degenerate IUPAC alphabet with 11 letters (ACGTRYSWSKN).

The IUPAC patterns can be transformed to PWMs based on the combined occurrences of all non-degenerated $K$-mers that match the degenerate IUPAC pattern in the input sequences. Alternatively, there is a faster approach based on the insight that if we allow any of the four nucleotides $a \in \{\text{A}, \text{C}, \text{G}, \text{T}\}$ at position $j$, the vast majority of motif matches will still be true positives due to the descriptive power of the other $K-1$ IUPAC letters. Therefore, we count the four nucleotides at motif position $j$ for matches to the pattern $y_{0:j-1}\text{N}y_{j+1:K-1}$ in which we replaced the $j$th IUPAC letter by an N:

$$p_{ja} = \frac{n(y_{0:j-1}\, a\, y_{j+1:K-1})}{n(y_{0:j-1}\, \text{N}\, y_{j+1:K-1})}\,, \tag{5}$$

where we have called $n(\mathbf{y})$ the number of occurrences of $K$-mer $\mathbf{y}$ in the input set. Note that these PWM probabilities can be computed solely from the $K$-mer counts in a time $O(W \times D)$ that is independent of the size of the input dataset $L_{\text{tot}}$, and only depends on the degeneracy $D = \left|\{\mathbf{x} \in \{\text{A}, \text{C}, \text{G}, \text{T}\}^W : \mathbf{x} \text{ matches } \mathbf{y}\}\right|$ of the motif $\mathbf{y}$, i.e., the number of different $K$-mers it matches.

We then refine the obtained PWMs by learning a multiple-occurrence-per-sequence model (MOPS) directly on the $K$-mer counts. The likelihood of a $K$-mer $\mathbf{x} \in \{\text{A}, \text{C}, \text{G}, \text{T}\}^K$ given a position weight matrix model with probabilities $\mathbf{p} = (p_j(\mathbf{A})$ is

$$\frac{p(\mathbf{x}|\mathbf{p}_{\text{motif}})}{p(\mathbf{x}|\mathbf{p}_{\text{bg}})} = \prod_{j=0}^{K-1} \frac{p_j(x_j)}{p_{\text{bg}}(x_j)}\,. \tag{6}$$

**Expectation step:** Compute the responsibilities $r(\mathbf{x})$, i.e., the probability that the factor will bind to $K$-mer $\mathbf{x}$.

$$r(\mathbf{x}) = \frac{p(\mathbf{x}|\mathbf{p}_{\text{motif}})/p(\mathbf{x}|\mathbf{p}_{\text{bg}})}{\sum_{\mathbf{x}' \in \{\text{A,C,G,T}\}^K} n(\mathbf{x}')p(\mathbf{x}'|\mathbf{p}_{\text{motif}})/p(\mathbf{x}'|\mathbf{p}_{\text{bg}})} \tag{7}$$

**Maximization step:** Update the probabilities of the position weight matrix model.

$$p_j(\mathbf{A}) = \sum_{\mathbf{x} \in \{\text{A,C,G,T}\}^K} I(x_j = a)\, n(\mathbf{x})\, r(\mathbf{x}) \tag{8}$$

By inserting the E-step equation into the M-step, we obtain

$$p_j^{(t)}(\mathbf{A}) \propto \sum_{\mathbf{x} \in \{\text{A,C,G,T}\}^K} I(x_j = a)\, n(\mathbf{x})\, \frac{p(\mathbf{x}|\mathbf{p}^{(t-1)}_{\text{motif}})}{p(\mathbf{x}|\mathbf{p}_{\text{bg}})} \tag{9}$$

and subsequent normalisation for each $j$ over $a \in \{\mathrm{A},\mathrm{C},\mathrm{G},\mathrm{T}\}$ yields the updated motif matrix probabilities.

To model saturation effects at the motifs with high affinities, we can use a saturation function that will limit the weight of the odds ratios to a maximum value $A$, e.g. $A = 1000$:

$$p_j^{(t)}(\mathbf{A}) \propto \sum_{\mathbf{x}\in\{\mathrm{A,C,G,T}\}^K} I(x_j = a)\, n(\mathbf{x}) \left( A^{-1} + \frac{p(\mathbf{x}|\mathbf{p}_{\mathrm{bg}})}{p(\mathbf{x}|\mathbf{p}_{\mathrm{motif}}^{(t-1)})} \right)^{-1} \tag{10}$$

In a thermodynamic interpretation, $A$ is the odds ratio of sites that have an occupancy of 50% at the assumed concentration of the transcription factor in the nucleus.

**Merging and extending PWMs.** We can reduce the redundancy of the PEnG!motif output and more importantly, generate more specific and sensitive motifs by merging sub-motifs that describe parts of the same underlying biological motif. For that, we first compute a list of pairwise similarity scores between all PWMs $\{p^{(1)}, \ldots, p^{(M)}\}$ with $P$-values above a user-specified cutoff obtained in the last step. Here, $p_{ja}^{(m)}$ is the probability of observing a nucleotide $a$ at the $j$'th position of that PWM. The similarity score $S(p^{(m)}, p^{(m')})$ is defined by the maximum similarity score $s(\cdot,\cdot)$ evaluated in the overlapping regions when the two patterns of length $l$ and $l'$ are shifted by $d = -2, -1, \ldots, l' - l + 2$ to each other:

$$S\left(p^{(m)}, p^{(m')}\right) = \max_{-2\le d\le l'-l+2} \left\{ s\left(p_{j_1:j_2}^{(m)}, p_{j'_1:j'_2}^{(m')}\right) \right\} . \tag{11}$$

The indices defining the overlap region in the two PWMs are $j_1 = \max\{0, d\}, j_2 = \min\{l - 1, l' - 1 + d\}$ and $j'_1 = \max\{0, -d\}, j'_2 = \min\{l' - 1, l - 1 - d\}$. The similarity score between the PWMs in the overlap region is computed using

$$s(p, p') = \frac{1}{2}\left(d(p, p^{(\mathrm{bg})}) + d(p', p^{(\mathrm{bg})})\right) - d(p, p'), \tag{12}$$

The distance $d(p, p')$ between two PWMs $p$ and $p'$ of length $l$ is the sum over the PWM columns of the relative entropies of each with their average distribution $\bar{p} := (p + p')/2$,

$$d(p, p') = \sum_{j=0}^{l-1} \left(H(p||\bar{p}) + H(p'||\bar{p})\right) = \sum_{j=0}^{l-1} \sum_{a\in\mathrm{A,C,G,T}} \left(p_{ja}\log_2 p_{ja} + p'_{ja}\log_2 p'_{ja} - 2\bar{p}_{ja}\log_2 \bar{p}_{ja}\right) . \tag{13}$$

The pair with the highest score will be merged using the positional offset $d$ that yielded the maximum similarity score. The pair of PWMs $(p^{(m)}, p^{(m')})$ has a score above a user-specified threshold ($0.4 \times W$ bits by default) are merged together using the positional offset $d$ that yielded the maximum similarity score. In the overlapping regions, the nucleotide probabilities of merged PWM will be the weighted sum of the nucleotide probabilities of the two merged PWMs, where the weights are the numbers of matches of the associated IUPAC patterns. The new weights of the columns of merged PWM will be the sum of these numbers of matches. In the non-overlapping regions, the probabilities and weights are simply copied over from the one PWM.

#### 1.2 Higher-order inhomogeneous Markov models

BaMMmotif [3] refines the pre-aligned short patterns or position-weight-matrices (PWMs) to higher-order Bayesian Markov models for the enriched motifs.

According to Boltzmann's law, the probability of a genomic site with sequence $\mathbf{x}$ to be bound by the transcription factor divided by the probability of $\mathbf{x}$ not to be bound is

$$\exp\left(-\frac{\Delta G(\mathbf{x})-\mu}{k_B T}\right) = \frac{p(\text{bound}|\mathbf{x})}{p(\text{not bound}|\mathbf{x})} = \frac{p(\text{bound}|\mathbf{x})}{1-p(\text{bound}|\mathbf{x})}, \tag{14}$$

with the chemical potential $\mu$ that depends on the factor concentration but not on $\mathbf{x}$. Solving for $p(\text{bound}|\mathbf{x})$ yields the well-known behaviour for saturated binding,

$$p(\text{bound}|\mathbf{x}) = \left(1+\exp\left(\frac{\Delta G(\mathbf{x})-\mu}{k_B T}\right)\right)^{-1}. \tag{15}$$

We parameterise the dependence of $\Delta G(\mathbf{x})$ on the binding site sequence $\mathbf{x}$ by a probability distribution $p_{\text{motif}}(\mathbf{x})$ which is defined by

$$p_{\text{motif}}(\mathbf{x})/p_{\text{bg}}(\mathbf{x}) \propto \exp(\Delta G(\mathbf{x})/k_B T). \tag{16}$$

The proportionality constant is determined by the normalization. Solving for $p_{\text{motif}}(\mathbf{x})$ and normalising yields

$$p_{\text{motif}}(\mathbf{x}) := \frac{p_{\text{bg}}(\mathbf{x})\exp(-\Delta G(\mathbf{x})/k_B T)}{\sum_{\mathbf{y}} p_{\text{bg}}(\mathbf{y})\exp(-\Delta G(\mathbf{y})/k_B T)}, \tag{17}$$

where the sum in the normalisation constant runs over all possible binding site sequences $\mathbf{y} \in \{\text{A},\text{C},\text{G},\text{T}\}^W$.

We derive a model for the Gibbs binding energy $\Delta G(\mathbf{x})$ for any potential binding site sequence $\mathbf{x} = x_{1:K} \in \{\text{A},\text{C},\text{G},\text{T}\}^K$ by computing a motif score $S(\mathbf{x})$:

$$S(\mathbf{x}) = -\frac{\Delta G(\mathbf{x})}{k_B T} + \text{const.} := \log\frac{p_{\text{motif}}(\mathbf{x})}{p_{\text{bg}}(\mathbf{x})} = \sum_{j=0}^{K-1}\log\frac{p_j^K(x_j|x_{j-K:j-1})}{p_{\text{bg}}^{K'}(x_j|x_{j-K':j-1})}. \tag{18}$$

where we model the background probability using a homogeneous Markov model of order $K'$:

$$p_{\text{bg}}(x_{i_0:i_1}) = \prod_{i=i_0}^{i_1} p_{\text{bg}}(x_i|x_{i-K':\,i-1}). \tag{19}$$

We model the motif using an inhomogeneous Markov model of order $K$:

$$p_{\text{motif}}(x_{0:K-1}) = \prod_{j=0}^{K-1} p_j(x_j|x_{j-K:\,j-1}). \tag{20}$$

We learn the parameters of the inhomogeneous Markov model by maximising the posterior probability. A natural prior is a product of Dirichlet distributions with pseudo-count parameters proportional to the lower-order model probabilities, with proportionality constants $\alpha_{kj}$ for $k = 1, \ldots, K$, whose size determines the strength of the prior. Maximizing the posterior probability yields

$$p_j^k(x_{k+1}|x_{1:k}) = \frac{n_j(x_{1:k+1}|\mathbf{r}) + \alpha_{kj} p_j^{k-1}(x_{k+1}|x_{2:k})}{n_{j-1}(x_{1:k}|\mathbf{r}) + \alpha_{kj}}. \tag{21}$$

#### 1.3 Masking in the motif refinement step

We train BaMMs using the expectation-maximization (EM) algorithm. In the E-step, we (re-) estimate the responsibilities $r$ for a motif to be present at position i of sequence n,

$$r_{ni} := p(z_n = i|\mathbf{x}_n, p^K_{\text{motif}}(\mathbf{x})) = \frac{p(\mathbf{x}_n|z_n = i, p^K_{\text{motif}}(\mathbf{x}))\, p(z_n = i)}{\sum_{i'=0}^{L_n-W+1} p(\mathbf{x}_n|z_n = i', p^K_{\text{motif}}(\mathbf{x}))\, p(z_n = i')} \tag{22}$$

In the M-step, we use the new $r_{ni}$ to update the model parameters $p_{\text{motif}}(\mathbf{x})^K$ for all orders $k = 0, ..., K$. This update equation looks exactly the same as the previous equation for known motifs locations, except that now the counts $n_j(x_{1:k+1})$ are interpreted as fractional counts computed according to

$$n_j(\mathbf{x}, x_{k+1}|\mathbf{r}) := \sum_{n=1}^{N} \sum_{i=1}^{L_n-W+1} r_{ni} I\left(x_{n,\, i+j-k\,:\,i+j} = (\mathbf{x}, x_{k+1})\right). \tag{23}$$

The indicator function $I$ returns 1 if the logical expression is true and 0 otherwise. The parameter updates are done for all orders from 0 to $K$.

Here we introduce a masking step between the E- and M-step by masking out the first N% of $r_{ni}$ after re-ranking increasingly (N is 90 by default) in the first iteration of the EM. By doing this, we learn the model only on the strong binding sites and thus eliminate the effect of unrelated motifs. We then iterate the EM algorithm until convergence.

#### 1.4 Optimization of order- and position-specific hyperparameters $\alpha$

In the previous version of BaMMmotif [3], the hyperparameters $\alpha_{kj}$ were empirically chosen. Here in this project, we try to learn the position-specific $\alpha_{kj}$ from the data.

We choose as prior on the hyperparameters $\alpha_{kj}$ (for $1 \leq k \leq K$) an inverse Gamma distribution with parameters 1 and $(\beta\gamma^k)$,

$$p(\alpha_{kj}|\beta, \gamma) = \frac{\beta\,\gamma^k}{\alpha_{kj}^2} e^{-\beta\,\gamma^k/\alpha_{kj}} \tag{24}$$

where $\beta \approx 5$ and $\gamma = 3$ corresponds roughly to the previous choice $\alpha_{kj} = \beta\,\gamma^k = 20 \times 3^{k-1}$ that worked for all of the datasets in the previous study [3].

According to Bayes' theorem, the conditional probability of $\boldsymbol{\alpha}$ given motif positions $\mathbf{z}$ can be written as:

$$
\begin{aligned}
p(\boldsymbol{\alpha}_k|\mathbf{X},\mathbf{z},p_{\text{motif}}^{k-1}) &\underset{\boldsymbol{\alpha}_k}{\propto} p(\mathbf{X}|\boldsymbol{\alpha},\mathbf{z},p_{\text{motif}}^{k-1})\, p(\boldsymbol{\alpha}|\mathbf{z},p_{\text{motif}}^{k-1}) \\
p(\boldsymbol{\alpha}_k|\mathbf{X},\mathbf{z},p_{\text{motif}}^{k-1}) &\underset{\boldsymbol{\alpha}_k}{\propto} p(\mathbf{X}|\boldsymbol{\alpha},\mathbf{z},p_{\text{motif}}^{k-1})\, p(\boldsymbol{\alpha}) \qquad (25)
\end{aligned}
$$

where

$$
\begin{aligned}
& p(\mathbf{X}|\mathbf{z},\boldsymbol{\alpha},p_{\text{motif}}^{k-1}) \\
& \propto \prod_{j=0}^{W-1}\prod_{\mathbf{y}} \frac{\Gamma(\alpha_{kj})}{\prod_a \Gamma(\alpha_{kj} v_j^*(a|\mathbf{y}'))} \frac{\prod_{a=1}^4 \Gamma(n_j^{\mathbf{z}}(\mathbf{y},a)+\alpha_{kj} v_j^*(a|\mathbf{y}'))}{\Gamma(n_{j-1}^{\mathbf{z}}(\mathbf{y})+\alpha_{kj})} \cancel{\prod_{a=1}^{4} \frac{1}{v_{\text{bg}}(a|\mathbf{y})^{n_j^{\mathbf{z}}(\mathbf{y},a)}}}. \qquad (26)
\end{aligned}
$$

Inserting (24) and (26) yields for the conditional probability

$$
\begin{aligned}
p(\boldsymbol{\alpha}_k|\mathbf{X},\mathbf{z},p_{\text{motif}}^{k-1}) &= \sum_{j=0}^{W-1} \left( \prod_{\mathbf{y}} \frac{\beta\,\gamma^k}{\alpha_{kj}^2} e^{-\frac{\beta\,\gamma^k}{\alpha_{kj}}} \frac{\Gamma(\alpha_{kj})}{\prod_a \Gamma(\alpha_{kj} v_j^*(a|\mathbf{y}'))} \frac{\prod_{a=1}^4 \Gamma(n_j^{\mathbf{z}}(\mathbf{y},a)+\alpha_{kj} v_j^*(a|\mathbf{y}'))}{\Gamma(n_{j-1}^{\mathbf{z}}(\mathbf{y})+\alpha_{kj})} \right) \\
&= \prod_{j=0}^{W-1} p(\alpha_{kj}|\mathbf{X},\mathbf{z},p_{\text{motif}}^{k-1})\,, \qquad (27)
\end{aligned}
$$

which factorizes over the $\alpha_{kj}$. We could therefore use Gibbs sampling to draw each new value of $\alpha_{kj}$ from its probability distribution independent of the others.

But for an efficient optimisation we need to reparameterise $\alpha_{kj}$ as

$$
\alpha_{kj} = e^{a_{kj}} \qquad (28)
$$

and sample $a_{kj}$ instead of $\alpha_{kj}$, because otherwise it would take too long to explore the entire probability distribution by small steps in $\alpha_{kj}$. If we went in steps of 0.5, for example, it would take almost 20000 directed steps to move from $\alpha_{kj} = 1$ to 10000. With steps of size 0.5, it would take only $2 log 20000 = 18.4$ directed steps to reach 10000. The probability density also needs to be transformed with the variable:

$$
\begin{aligned}
p(a_{kj}|\mathbf{X},\mathbf{z},p_{\text{motif}}^{k-1}) &= \left|\frac{d\,\alpha_{kj}}{d\,a_{kj}}\right| p(\alpha_{kj}|\mathbf{X},\mathbf{z},p_{\text{motif}}^{k-1}) \qquad (29) \\
&= \alpha_{kj}\, p(\alpha_{kj}|\mathbf{X},\mathbf{z},p_{\text{motif}}^{k-1}) \qquad (30)
\end{aligned}
$$

The log conditional probability for $a_{kl}$ is

$$
\begin{aligned}
& \log p(a_{kl}|\mathbf{X},\mathbf{z},p_{\text{motif}}^{k-1}) = \text{const.} - \log\alpha_{kj} - \beta\,\gamma^k/\alpha_{kj} + 4^k \log\Gamma(\alpha_{kj}) \qquad (31) \\
& + \sum_{\mathbf{y}=y_{1:k}} \left( \sum_{a=1}^{4} \left[ \log\Gamma(n_j^{\mathbf{z}}(\mathbf{y},a)+\alpha_{kj} p_{\text{motif},j}^{k-1}(a|\mathbf{y}')) - \log\Gamma(\alpha_{kj} v_j^{k-1}(a|\mathbf{y}')) \right] - \log\Gamma(n_{j-1}^{\mathbf{z}}(\mathbf{y})+\alpha_{kj}) \right)
\end{aligned}
$$

We can sample from this distribution using the Metropolis-Hastings algorithm. We draw a new $a_{kl}^{\text{try}} \sim \mathcal{N}(a_{kl}, 1)$ and accept this trial sample with a probability

$$\frac{p(a_{kl}^{\text{try}}|\mathbf{X}, \mathbf{z}, p_{\text{motif}}^{k-1})}{p(a_{kl}|\mathbf{X}, \mathbf{z}, p_{\text{motif}}^{k-1})} \text{ if } p(a_{kl}^{\text{try}}|\mathbf{X}, \mathbf{z}, p_{\text{motif}}^{k-1}) < p(a_{kl}|\mathbf{X}, \mathbf{z}, p_{\text{motif}}^{k-1})$$
$$1 \text{ if otherwise .} \quad (32)$$

Because it is fast to sample $a_{kl}$ in this way, we draw 10 or times in a row and only take record the last accepted sample of $a_{kl}$. This 10-fold repetition ensures that we can explore almost the entire range of relevant values of $a_{kl}$ within these 10 steps.

At the start of the sampling, the $a_{kj}$ will move in the direction of the medians of their probability distribution in relatively directed steps until the changes to the $a_{kj}$ become non-directional and begin to fluctuate. We can then fix the $a_{kj}$ to the average of the last 20 or so samples and perform a few (e.g. 5) iterations of the EM algorithm (described in section 1.2) to find the optimum model parameters $v_j^K(a|\mathbf{y})$ given the fixed $a_{kj}$.

#### 1.5 Learning positional preferences of motifs

##### Thermodynamic treatment of positional preference

We proceed analogously to section 1.2 but introduce a positional preference as an additive term $\Delta G_i$ in the binding energy. The probability of a factor to bind a binding site consisting of W nucleotides between $i$ and $i + W - 1$ in a sequence $\mathbf{x} = x_{1:L}$ then becomes

$$p(\text{factor bound at position } i|\mathbf{x}) = \left(1 + \exp\left(\frac{\Delta G(x_{i:i+W-1}) + \Delta G_i - \mu}{k_B T}\right)\right)^{-1} . \quad (33)$$

We define $p_{\text{motif}}(x_{0:W-1})$ as in eq. (17) and we further define a positional distribution

$$p(z = i|\text{factor bound to } \mathbf{x}) = \frac{\exp(-\Delta G_i/k_B T)}{\sum_{i'=1}^{L} \exp(-\Delta G_{i'}/k_B T)} . \quad (34)$$

We abbreviate the denominator as const. gives

$$-\frac{\Delta G_i}{k_B T} + \text{const.} = \log p(z = i|\text{factor bound to } \mathbf{x}) =: s_i. \quad (35)$$

Once we know $p_{\text{motif}}(\cdot)$ and $p(z = i|\text{factor bound to } \mathbf{x})$, we can compute $S(x_{i:i+W-1})$ and $s_i$ and the relative binding strength $(\Delta G(x_{i:i+W-1}) + \Delta G_i)/k_B T$ for any potential binding site position $i$ in any sequence $\mathbf{x} = (x_1 \dots x_L)$.

If we again assume to be in a regime of unsaturated binding, $p(\text{bound}|\mathbf{x}) \lesssim 0.1$ we can approximate the probability $p(\mathbf{x}_n|\text{bound}, p_{\text{motif}}^k)$ for pulling out a sequence $\mathbf{x}_n$ from an underlying distribution of possible sequences $p_{\text{bg}}(\mathbf{x})$ as

$$\begin{aligned}
p(\mathbf{x}_n|\text{bound}, p^k_{\text{motif}}) &\propto p(\text{factor bound}|\mathbf{x}_n, p^k_{\text{motif}})\, p_{\text{bg}}(\mathbf{x}_n) \\
&= p_{\text{bg}}(\mathbf{x}_n) \sum_{i=1}^{L-W+1} p(\text{factor bound at } i|\mathbf{x}_n, p^k_{\text{motif}}) \\
&= p_{\text{bg}}(\mathbf{x}_n) \sum_{i=1}^{L-W+1} \left(1 + \exp\left(\frac{\Delta G(x_{i:i+W-1}) + \Delta G_i - \mu}{k_B T}\right)\right)^{-1} \\
&\approx p_{\text{bg}}(\mathbf{x}_n) \sum_{i=1}^{L-W+1} \exp\left(-\frac{\Delta G(x_{i:i+W-1}) + \Delta G_i - \mu}{k_B T}\right) \\
&\propto p_{\text{bg}}(\mathbf{x}_n) \sum_{i=1}^{L-W+1} \exp\left(S(x_{i:i+W-1}) + s_i\right). \qquad (36)
\end{aligned}$$

To find the model parameters $\boldsymbol{\theta}$ consisting of $\mathbf{s} = (s_1, \ldots, s_{L-W+1})$ and of $p^k_{\text{motif}}$ specifying $p_{\text{motif}}(\cdot)$, we need to optimise the log likelihood function of these parameters:

$$LL(\boldsymbol{\theta}) = \sum_{n=1}^{N} \log p(\mathbf{x}_n|\text{bound}, p^k_{\text{motif}}, \mathbf{s}) \qquad (37)$$

##### Flat Bayesian prior on positional preference

Let us define parameters $\boldsymbol{\pi}$ with $\pi_i = p(z{=}i|z_i{\neq}0) = e^{s_i}$ the probability of a motif to start at position $i$ of a sequence. The M-step will then be given again by equation (22) but this time using the positional preferences $\pi_i$ instead of the flat positional distribution. We will use a flat prior distribution,

$$p(\boldsymbol{\pi}|\beta) = \text{Dir}(\boldsymbol{\pi}|\beta\mathbb{1}), \qquad (38)$$

and we will choose a value around $\beta = 2 \ldots 10$.

The auxiliary function becomes

$$
\begin{aligned}
&Q(p^k_{\text{motif}}, \boldsymbol{\alpha}, q|\mathbf{r}, p^{k-1}_{\text{motif}}) \\
&= \sum_{n=1}^{N} \left[ \sum_{i=0}^{L_n-W+1} r_{ni} \log \left( p(\mathbf{x}_n|z_n=i, p^k_{\text{motif}})\, p(z_n=i|q) \right) \right] + \log p(p^k_{\text{motif}}|p^{k-1}_{\text{motif}}, \boldsymbol{\alpha}) + \log p(\boldsymbol{\pi}|\beta) \\
&= \sum_{n=1}^{N} \sum_{i=0}^{L_n-W+1} r_{ni} \log p(\mathbf{x}_n|z_n=i, p^k_{\text{motif}}) + \log p(p^k_{\text{motif}}|p^{k-1}_{\text{motif}}, \boldsymbol{\alpha}) \\
&\quad + \sum_{n=1}^{N} \left( r_{n,0} \log(1-q) + \sum_{i=1}^{L_n-W+1} r_{ni} \log(q\pi_i) \right) + \log \text{Dir}(\boldsymbol{\pi}|\beta\mathbb{1}) \\
&= \sum_{n=1}^{N} \sum_{i=0}^{L_n-W+1} r_{ni} \log p(\mathbf{x}_n|z_n=i, p^k_{\text{motif}}) + \log p(p^k_{\text{motif}}|p^{k-1}_{\text{motif}}, \boldsymbol{\alpha}) \qquad (39) \\
&\quad + \sum_{n=1}^{N} \left( r_{n,0} \log(1-q) + (1-r_{n,0}) \log q + \sum_{i=1}^{L_n-W+1} r_{ni} \log \pi_i \right) + \sum_{i=1}^{L_n-W+1} (\beta-1) \log \pi_i .
\end{aligned}
$$

We use the method or Lagrange multipliers again to find the optimum of $Q(p^k_{\text{motif}}, \boldsymbol{\alpha}, q|\mathbf{r}, p^{k-1}_{\text{motif}})$ under the constraint $\sum_{i=1}^{L-W+1} \pi_i = 1$:

$$
\frac{\partial}{\partial \pi_i} \left( Q(p^k_{\text{motif}}, \boldsymbol{\alpha}, q|\mathbf{r}, p^{k-1}_{\text{motif}}) - \lambda \left( \sum_{i=1}^{L-W+1} \pi_i - 1 \right) \right) = \sum_{n=1}^{N} \frac{r_{ni}}{\pi_i} + \frac{\beta-1}{\pi_i} - \lambda = 0 \qquad (40)
$$

Solving for $\pi_i$, normalising the distribution and defining $N_i := \sum_{n=1}^{N} r_{ni}$ yields

$$
\boxed{\pi_i = \frac{N_i + \beta - 1}{N + (L-W+1)(\beta-1)}\,.} \qquad (41)
$$

#### Prior penalising jumps in the positional preference profile

For many applications it might be more appropriate to limit the complexity of the positional preference profile by imposing a smoothness on the $p(z=i)$. For example, transcription factor binding sites will be more frequent near the center of ChIP-seq peaks than farther away; factors bind more strongly to the outer parts of probes on protein binding microarrays than to the parts near the glass slide; transcription factors in HT-SELEX experiments might prefer the center of probes over the ends. In the following we assume that all training and test sequences have the same length $L$.

Because the smoothness prior couples neighbouring positional probabilities with each other, there is no closed-form solution for the parameters anymore. We have to use a gradient-based optimisation such as conjugate gradients to minimise $Q$ with respect to the positional parameters. We therefore parameterise the positional distribution in such a way that the normalisation condition $\sum_i \pi_i = 1$ and the limits $0 \leq \pi_i \leq 1$ automatically hold true during the numerical optimisation,

$$
p(z_n\!=\!i|z_n\!\neq\!0) = \frac{e^{s_i}}{\sum_{i'=1}^{L-W+1} e^{s_{i'}}}. \qquad (42)
$$

We impose a smoothness prior on the $\pi_i$, that encourages the point-wise estimated first derivative to stay small,

$$p(\boldsymbol{\pi}|\beta) = \prod_{i=2}^{L-W+1} \mathcal{N}\left(s_i - s_{i-1}|\,0, \beta^{-1}\right) \,, \tag{43}$$

with precision (= inverse variance) $\beta$.

With this prior, the auxiliary function becomes

$$\begin{aligned} Q(p^k_{\text{motif}}, \boldsymbol{\alpha}, q, \boldsymbol{\pi}|\mathbf{r}, p^{k-1}_{\text{motif}}) = &\sum_{n=1}^{N}\sum_{i=0}^{L-W+1} r_{ni} \log p(\mathbf{x}_n|z_n = i, p^k_{\text{motif}}) + \log p(p^k_{\text{motif}}|p^{k-1}_{\text{motif}}, \boldsymbol{\alpha}) \\ &+ \sum_{n=1}^{N}\left( r_{n,0}\log(1-q) + (1-r_{n,0})\log q + \sum_{i=1}^{L-W+1} r_{ni}\left( s_i - \log\left(\sum_{i'} e^{s_{i'}}\right)\right)\right) \\ &- \frac{\beta}{2}\sum_{i=2}^{L-W+1} (s_i - s_{i-1})^2 + \frac{L-W}{2}\log\beta + \text{const.} \end{aligned} \tag{44}$$

The partial derivatives of $Q(p^k_{\text{motif}}, \boldsymbol{\alpha}, q, \boldsymbol{\pi}|\mathbf{r}, p^{k-1}_{\text{motif}})$ are

$$\begin{aligned} \frac{\partial}{\partial s_i} Q(p^k_{\text{motif}}, \boldsymbol{\alpha}, q, \boldsymbol{\pi}|\mathbf{r}, p^{k-1}_{\text{motif}}) = &\sum_{n=1}^{N} r_{ni} - \sum_{n=1}^{N}\sum_{i'=1}^{L-W+1} r_{ni'} \frac{e^{s_i}}{\sum_{i''} e^{s_{i''}}} \\ &- \beta\,(s_i - s_{i-1})\, I(2 \le i \le L-W+1) \\ &+ \beta\,(s_{i+1} - s_i)\, I(1 \le i \le L-W) \end{aligned} \tag{45}$$

and

$$\boxed{\frac{\partial}{\partial s_i} Q(p^k_{\text{motif}}, \boldsymbol{\alpha}, q, \boldsymbol{\pi}|\mathbf{r}, p^{k-1}_{\text{motif}}) = N_i - (N - N_0)\, p(z{=}i|z{\neq}0) - (\beta \mathbf{A}\boldsymbol{s})_i}$$

with the abbreviations $N_0 := \sum_{n=1}^{N} r_{n,0}$ and

$$\mathbf{A} := \begin{pmatrix} 1 & -1 & 0 & 0 & \cdots & \cdots & \cdots & 0 \\ -1 & 2 & -1 & 0 & \ddots & \ddots & \ddots & \vdots \\ 0 & -1 & 2 & -1 & \ddots & \ddots & \ddots & \vdots \\ 0 & 0 & -1 & 2 & \ddots & \ddots & \ddots & \vdots \\ \vdots & \ddots & \ddots & \ddots & \ddots & \ddots & 0 & 0 \\ \vdots & \ddots & \ddots & \ddots & \ddots & 2 & -1 & 0 \\ \vdots & \ddots & \ddots & \ddots & 0 & -1 & 2 & -1 \\ 0 & \cdots & \cdots & \cdots & 0 & 0 & -1 & 1 \end{pmatrix} . \tag{46}$$

The partial derivative will adjust $s_i$ such that $p(z{=}i|z{\neq}0) = e^{s_i}/\sum_{i'} e^{s_{i'}}$ equals $N_i/(N - N_0)$ plus a smoothness correction $\mathbf{A}\boldsymbol{s}$ that will pull $s_i$ up or down in order to minimise the estimator of the second derivative of the profile at position $i$. We run a few iterations of conjugate gradients (e.g. 5 to 10) during each EM step to learn the positional preferences.

**Learning the optimal smoothness parameter $\beta$ from the data.** We can regard $Q$ also as a function of $\beta$,

$$Q(p^k_{\text{motif}}, \boldsymbol{\alpha}, q, \boldsymbol{\pi}, \beta | \mathbf{r}, p^{k-1}_{\text{motif}}) = -\frac{\beta}{2} \sum_{i=2}^{L-W+1} (\pi_i - \pi_{i-1})^2 + \frac{L-W}{2} \log \beta + \text{const}_\beta \,, \tag{47}$$

and optimise is with respect to $\beta$:

$$0 = \frac{\partial}{\partial \beta} Q(p^k_{\text{motif}}, \boldsymbol{\alpha}, q, \boldsymbol{\pi}, \beta | \mathbf{r}, p^{k-1}_{\text{motif}}) = -\frac{1}{2} \sum_{i=2}^{L-W+1} (s_i - s_{i-1})^2 + \frac{L-W}{2\beta} \tag{48}$$

and therefore

$$\boxed{\beta = \left( \frac{1}{L-W} \sum_{i=2}^{L-W+1} (s_i - s_{i-1})^2 \right)^{-1}} \tag{49}$$

Instead of optimising $\beta$, we can again interpret $Q$ as the likelihood of an ensemble of fractional motif instances with weights $r_{ni}$ and compute the expectation value of $\beta$. If we assume a uniform prior on $\beta$, $p(\beta) = \text{const}$, the posterior distribution of $\beta$ is proportional to the likelihood. We note that the functional form of $Q(\beta)$ is that of a Gamma distribution, $Q(\beta) = \log \text{Ga}(\beta | a, b) + \text{const} = (a-1) \log \beta - b\beta + \text{const}$, with $a - 1 = (L-W)/2$ and $b = (1/2) \sum_i (s_i - s_{i-1})^2$. Since the expectation value of a Gamma distribution is $a/b$, we can conclude for $\beta$

$$\boxed{\mathbb{E}[\beta] = \left( \frac{1}{L-W+2} \sum_{i=2}^{L-W+1} (s_i - s_{i-1})^2 \right)^{-1} .} \tag{50}$$

We can then update $\beta$ by its expectation value instead of the mode of $Q(\beta)$. Alternatively, we could sample $\beta$ from the Gamma distribution $\text{Ga}(\beta | (L-W+2)/2, (1/2) \sum_i (s_i - s_{i-1})^2)$.

#### Prior penalising kinks in the positional preference profile

For various applications such as PBMs and HT-SELEC, we might be interested in more smooth positional preferences. In these cases, it might be better to use a smoothness prior on the $\pi_i$ that encourages the point wise estimated *third* derivative to stay small,

$$p(\boldsymbol{\pi} | \beta) = \prod_{i=2}^{L-W} \mathcal{N} \left( s_i - \frac{s_{i-1} + s_{i+1}}{2} \middle| 0, \beta^{-1} \right) , \tag{51}$$

with precision (= inverse variance) $\beta$. With this prior, the auxiliary function becomes

$$\begin{aligned} Q(p^k_{\text{motif}}, \boldsymbol{\alpha}, q, \boldsymbol{\pi} | \mathbf{r}, p^{k-1}_{\text{motif}}) = & \sum_{n=1}^{N} \sum_{i=0}^{L-W+1} r_{ni} \log p(\mathbf{x}_n | z_n = i, p^k_{\text{motif}}) + \log p(p^k_{\text{motif}} | p^{k-1}_{\text{motif}}, \boldsymbol{\alpha}) \\ & + \sum_{n=1}^{N} \left( r_{n,0} \log(1-q) + (1-r_{n,0}) \log q + \sum_{i=1}^{L-W+1} r_{ni} \left( s_i - \log \left( \sum_{i'} e^{s_{i'}} \right) \right) \right) \\ & - \frac{\beta}{2} \sum_{i=2}^{L-W} \left( s_i - \frac{s_{i-1} + s_{i+1}}{2} \right)^2 + \frac{L-W-1}{2} \log \beta + \text{const}. \end{aligned} \tag{52}$$

The partial derivatives of $Q(p^k_{\text{motif}}, \boldsymbol{\alpha}, q, \boldsymbol{\pi} | \mathbf{r}, p^{k-1}_{\text{motif}})$ are

$$\begin{aligned}\frac{\partial}{\partial s_i} Q(p^k_{\text{motif}}, \boldsymbol{\alpha}, q, \boldsymbol{\pi} | \mathbf{r}, p^{k-1}_{\text{motif}}) = \sum_{n=1}^{N} r_{ni} - \sum_{n=1}^{N} \sum_{i'=1}^{L-W+1} r_{ni'} \frac{e^{s_i}}{\sum_{i''} e^{s_{i''}}} \\ + \frac{\beta}{2} \left( s_{i-1} - \frac{s_{i-2} + s_i}{2} \right) I(3 \le i \le L-W+1) \\ - \beta \left( s_i - \frac{s_{i-1} + s_{i+1}}{2} \right) I(2 \le i \le L-W) \\ + \frac{\beta}{2} \left( s_{i+1} - \frac{s_i + s_{i+2}}{2} \right) I(1 \le i \le L-W-1) \end{aligned} \tag{53}$$

and

$$\boxed{\frac{\partial}{\partial s_i} Q(p^k_{\text{motif}}, \boldsymbol{\alpha}, q, \boldsymbol{\pi} | \mathbf{r}, p^{k-1}_{\text{motif}}) = N_i - (N - N_0)\, p(z{=}i | z{\neq}0) - \frac{\beta}{4} (\mathbf{B}\boldsymbol{s})_i}$$

with the abbreviations $N_0 := \sum_{n=1}^{N} r_{n,0}$ and

$$\mathbf{B} := \begin{pmatrix} 1 & -2 & 1 & 0 & 0 & \cdots & \cdots & \cdots & 0 \\ -2 & 5 & -4 & 1 & 0 & \ddots & \ddots & \ddots & \vdots \\ 1 & -4 & 6 & -4 & 1 & \ddots & \ddots & \ddots & \vdots \\ 0 & 1 & -4 & 6 & -4 & \ddots & \ddots & \ddots & \vdots \\ 0 & 0 & 1 & -4 & 6 & \ddots & \ddots & \ddots & \vdots \\ \vdots & \ddots & \ddots & \ddots & \ddots & \ddots & \ddots & 1 & 0 \\ \vdots & \ddots & \ddots & \ddots & \ddots & \ddots & 6 & -4 & 1 \\ \vdots & \ddots & \ddots & \ddots & \ddots & 1 & -4 & 5 & -2 \\ 0 & \cdots & \cdots & \cdots & \cdots & 0 & 1 & -2 & 1 \end{pmatrix}. \tag{54}$$

The partial derivative will adjust $s_i$ such that $p(z{=}i|z{\neq}0) = e^{s_i} / \sum_{i'} e^{s_{i'}}$ equals $N_i/(N - N_0)$ plus a smoothness correction $\mathbf{B}\boldsymbol{s}$ that will pull $s_i$ up or down in order to minimise the estimator of the third derivative of the profile at position $i$.

**Learning the optimal smoothness parameter $\beta$ from the data.** Analogously to the previous smoothness prior, we can learn $\beta$ from the data using the update

$$\boxed{\beta = \left( \frac{1}{L-W-1} \sum_{i=2}^{L-W} \left( s_i - \frac{s_{i-1} + s_{i+1}}{2} \right)^2 \right)^{-1}} \tag{55}$$

or

$$\boxed{\beta = \left( \frac{1}{L-W+1} \sum_{i=2}^{L-W} \left( s_i - \frac{s_{i-1} + s_{i+1}}{2} \right)^2 \right)^{-1}.} \tag{56}$$

#### 1.6 Scanning sequences for motif occurrences

To obtain the motif occurrences from the sequences, given a known or learned motif, we developed a motif scanning tool BaMMScan to evaluate the possible motif occurrences on the input sequences. The motif score $s_i(x_{1:K})$ is calculated for each position $i$ on every sequence $x$ for the order $K$. A background score distribution is created by generating $M$-fold background sequences from a second-order homogeneous Markov model from input set ($M$ can be 10). We sort the list of $N^+ + N^-$ positive- and negative-set scores jointly in descending order. We denote the cumulative number of scores from the negative set up to rank $l$ in this list by $\mathrm{FP}_l$ and then compute the P-value of entry $l$ with score $S_l$ in that list by

$$P\text{-value}(S_l) = \frac{1}{N^-}\left(\mathrm{FP}_l + \frac{S_l^{\text{higher}} - S_l}{S_l^{\text{higher}} - S_l^{\text{lower}} + \epsilon}\right). \tag{57}$$

and the $E$-values are obtained simply as

$$E\text{-value} = N^+ \times P\text{-value}. \tag{58}$$

The motif occurrences with a $P$-value smaller than certain cutoff (e.g. $1e^{-4}$) are reported.

#### 1.7 Evaluation criteria using the average recall (AvRec) score

To assess the predictive performance of the motif finders, we first defined an average recall (AvRec) score (details also described in [4]). The AvRec score represents the averaged recall over the range of precision from 0 to 1. The advantage of AvRec score over commonly used $p$-value is that it covers the most relevant range of False-discovery-rates (FDR) in practical applications and allows the user to intuitively estimate the motif performance in her particular application.

We obtain a $p$-value for each sequence by

$$p\text{-value}_l = \frac{\mathrm{FP}_l + 0.5}{\mathrm{N}^- + 1} \tag{59}$$

After having a $p$-value for every motif occurrence (as described in eq.57), we obtain a list of corresponding local FDR values and an estimate of the weight of the null component $\eta_0$ by applying fdrtool [5] on the $p$-value distribution (Figure S 13A). We then calculate FDR and recall for each entry by

$$\mathrm{FDR}_l = \frac{\mathrm{FP}_l}{\mathrm{FP}_l + \mathrm{TP}_l} \tag{60}$$

$$\mathrm{recall}_l = (1 - \mathrm{FDR}_l)\frac{l}{(1-\eta_0)N} \tag{61}$$

The ratio between true positive (TP) and false positive (FP) is calculated by

$$R_{l[\mathrm{TP/FP}]} = \frac{1 - \mathrm{FDR}_l}{\mathrm{FDR}_l} \times M \tag{62}$$

with $M$ as the ratio between negative and positive sequences.

We visualize the characteristics by plotting the TP/FP ratio $R_{\mathrm{TP/FP}}$ on the y-axis against the recall on the x-axis, and define the calculated area-under-the-curve as the AvRec score for motif evaluation (Figure S 13B).

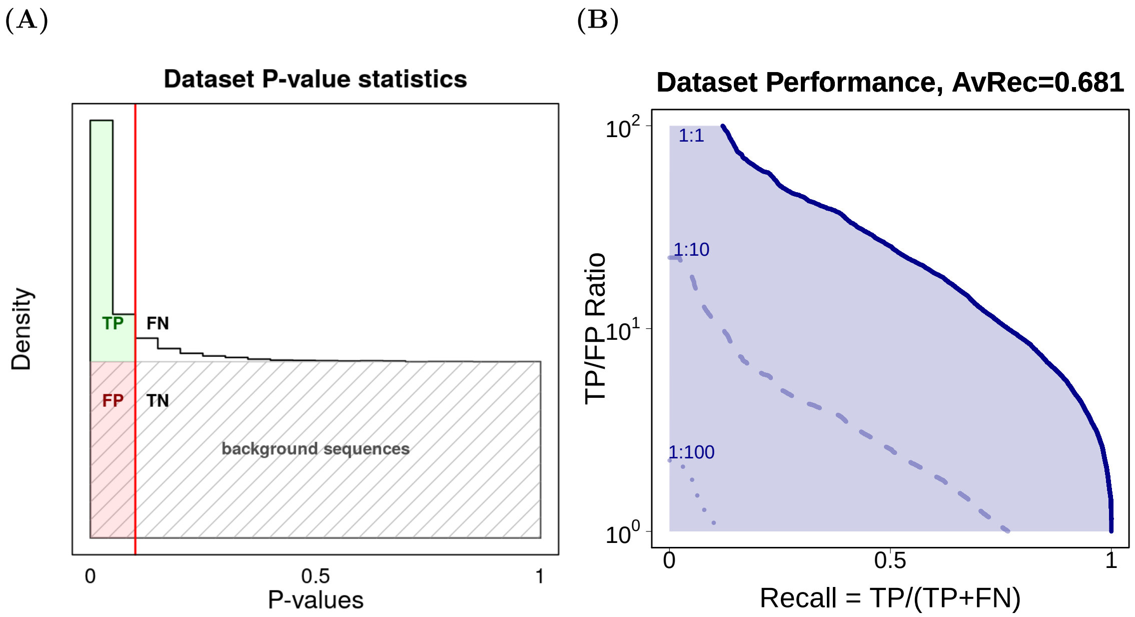


**Figure S 13. Schemes of motif assessment on sequences.** (**A**) We calculate p-values for the most likely positions on both positive and background sequences, given the motif and a second-order background model learned from the sequences. We plot the density of the p-values and choose a cutoff at 0.1 (the solid red line). The background sequences are mapped in the grey shadow, given the ratio between the background and positive sequences. The true positives (TP, in green), false negatives (FN, in white), false positives (FP, in red) and true negatives (TN) are visualized on the plot. (**B**) For each p-value for positive sequences, we calculate the recall and the ratio between TP and FP, and then plot the recall against the ratio of TP/FP. The solid dark blue line represent for the scenario when the ratio between positive and background sequences is 1:1, the dash lines under it are for the cases when the ratio is 1:10 and 1:100, respectively. An average recall (AvRec) score is calculated as the area under the curve for the 1:1 ratio scenario, and used as a measurement for motif quality on the positive sequence.

### 2 Datasets used for the benchmark

#### 2.1 ENCODE database

We evaluated the performance of the selected algorithms on human ChIP-seq datasets from the ENCODE portal [6] till March 2012. In total, there are 435 datasets for 93 distinct transcription factors. The top 5000 peak regions, sorted by their signal value, were selected for each dataset. If fewer than 5000 peaks were contained in a dataset, all peaks were chosen. Positive sequences were extracted ±104 bp around the peak summits. Background sequences were sampled by trimer frequencies from positive sequences, with the same length as positive sequences and 10 times the amount of positive sequences. 8 datasets were excluded from all the results because diChIPMunk failed to learn models within 3 hours.

##### 2.2 HT-SELEX datasets

For HT-SELEX data, we downloaded 164 datasets with 200 bp-long oligomers from Zhu et al. [7], which are deposited in the European Nucleotide Archive (ENA) under accession PRJEB22684. Each dataset represents one non-redundant human transcription factor. For each dataset, we selected the top 5000 sequences from the 4th cycle without any sorting. Background sequences are sampled in the same way as described previously.

##### 2.3 GTRD database

For the GTRD database, we obtained 405 *in vivo* datasets for 405 non-redundant human transcription factors from Yevshin et al. [1]. The top 5000 peak regions are selected after sorting by q-values. Positive sequences are extracted ±100 bp around the peak summits. Background sequences are sampled in the same way as described previously.

##### 2.4 MITOMI datasets

MITOMI is a microfluidics-based approach for *de novo* discovery and quantitative biophysical characterization of DNA target sequences [8]. We downloaded the MITOMI data for 28 *Saccharomyces cerevisiae* transcription factors under the accession GPL10817. The 3 bp and 15 bp long adapters on both ends are truncated. We then downloaded yeast GTRD datasets for 8 transcription factors [1] for the motif discovery.

##### 2.5 Cross-platform datasets

Out of 435 ENCODE datasets for 93 TFs and 164 HT-SELEX datasets for 164 non-redundant TFs, there are 66 TFs which have both *in vivo* and *in vitro* datasets. Out of 66 TFs, most of them have very low AvRec scores when performing the cross-platform validations. We investigated into details and found out that for most of them, the learned motifs were very distinct from the two platforms. This result confirms that TFs can bind to different motifs when experimenting either *in vivo* or *in vitro*. For the left 16 paired tests, they are motifs for 4 TFs, namely CEBPB, POU2F2, ELF1, and FOXA2, which were used in our benchmark.

#### 3 Motif finders used in the benchmark

The source code is available for command-line versions of PEnGmotif and BaMMmotif2 and supported on Linux and MacOS:

##### 3.1 PEnGmotif

PEnGmotif repository: github.com/soedinglab/PEnG-motif. For this study, we used parameters `--optimization_score MUTUAL_INFO -w 8 --threads 4`. The output is in MEME-like format. The motifs are sorted by their AvRec scores, and the best one was taken for the benchmark.

##### 3.2 BaMMmotif2

BaMMmotif2 repository: github.com/soedinglab/BaMMmotif2. For this study, we seeded with the PWMs discovered by PEnGmotif and used parameters `--EM -k [k] --advanceEM --extend 2 2` for further optimization. `[k]` is chosen as 1 and 5 for the benchmark for this study. The output format is defined as BaMM format with extensions like `.ihbcp` and `.hbcp`.

##### 3.3 BaMMmotif

BaMMmotif repository: github.com/soedinglab/BaMMmotif. For this study, we seeded with PWMs by triggering XXmotif internally and used parameters `--reverseComp --XX-localization --XX-localizationRanking --XX-K 2 --maxPValue 0.05 --maxPWMs 3 --extend 2 2` for further optimization. The output format is defined as BaMM format with extensions like `.ihbcp` and `.hbcp`.

##### 3.4 CisFinder

CisFinder was installed from https://lgsun.grc.nia.nih.gov/CisFinder/download.html. We ran patternFind for identifying motifs, patternCluster for clustering motifs, and patternTest for improving motifs. Default parameters were applied. The discovered motifs were converted to MEME-like output format and re-ranked by our motif sorting script, and only the best motif was taken for the benchmarks.

##### 3.5 MEME

MEME version 5.1.1 was installed and applied with parameters `-dna -mod zoops -nmotifs 3 -revcomp -p 4 -V 2`. Maximum 3 motifs were saved in the output, and the best one according to the AvRec score was taken for the benchmarks.

##### 3.6 ChIPMunk

ChIPMunk version v8 was downloaded and applied with parameters `ru.autosome.ChIPMunk 8 12 yes 1.0 100 10 1 4`. The discovered motifs were converted to MEME-like output format.

#### 3.7 diChIPMunk

diChIPMunk was implemented in the same package as ChIPMunk. We ran it with parameters `ru.autosome.di.ChIPMunk 8 12 yes 1.0 200 20 1 4`. The discovered motifs were converted to BaMM-like output format for further comparison.

#### 3.8 InMoDe

InMoDe was downloaded from http://www.jstacs.de/index.php/InMoDe. We applied the module `flexible`, which allows us to customize the learning task. The discovered motifs were converted to BaMM-like output format for further comparison.
